# Murder in cold blood? Forensic and bioarchaeological identification of the skeletal remains of Béla, Duke of Macsó (c. 1245–1272)

**DOI:** 10.1101/2025.07.25.666716

**Authors:** Tamás Hajdu, Noémi Borbély, Zsolt Bernert, Ágota Buzár, Tamás Szeniczey, István Major, Claudio Cavazzuti, Mihály Molnár, Anikó Horváth, László Palcsu, Barna Árpád Kelentey, János Angyal, Balázs Gusztáv Mende, Kristóf Jakab, Zsuzsa Lisztes-Szabó, Ágoston Takács, Olivia Cheronet, Ron Pinhasi, David Emil Reich, Martin Trautmann, Anna Szécsényi-Nagy

**Affiliations:** Department of Biological Anthropology, Faculty of Science, Eötvös Loránd University, Budapest, Hungary; Institute of Archaeogenomics, Research Centre for the Humanities, HUN-REN; Department of Anthropology, Hungarian Natural History Museum, Budapest, Hungary; Department of Anthropology, Hungarian Natural History Museum, Budapest, Hungary.; Isotope Climatology and Environmental Research Centre, HUN-REN Institute for Nuclear Research, Debrecen, Hungary; Department of History and Culture, Alma Mater Studiorum, University of Bologna, Bologna, Italy; Department of Operative Dentistry and Endodontics, Faculty of Dentistry, University of Debrecen, Debrecen, Hungary; Department of Periodontology, Faculty of Dentistry, University of Debrecen, Debrecen, Hungary; Institute of Archaeogenomics, HUN-REN Research Centre for the Humanities, Budapest, Hungary; Medieval Department, Castle Museum - Budapest History Museum, Budapest, Hungary; University of Vienna; Department of Genetics, Harvard Medical School, Boston, USA; Department of Cultures/Archaeology, University of Helsinki, Helsinki, Finland

**Keywords:** Forensic identification, Ancient DNA (aDNA), Bioarchaeology, Identity by descent (IBD), Medieval royal remains, Perimortem trauma analysis, Árpád, Rurikid dynasties

## Abstract

In 1915, the remains of a male were discovered in a 13th-century monastery on Margaret Island, Budapest. Historical context suggested that the remains may belong to Duke Béla of Macsó (c. 1245–1272), grandson of King Béla IV of Hungary (House of Árpád) and son of Duke Rostislav (Rurik dynasty). We applied a complex approach to identify the individual and reconstruct the circumstances of his death. Radiocarbon dating, when adjusted for freshwater reservoir effects linked to a high-protein diet, placed the burial in the mid-13th century. Skeletal features corresponded to a young adult male. Stable isotope and dental calculus analyses indicated a high-status diet rich in animal proteins and C3 cereals. Ancient DNA confirmed descent from King Béla III (Árpád dynasty) and Y-chromosomal affiliation with the Rurikid lineage. Forensic evidence revealed 26 perimortem injuries, suggesting a coordinated, premeditated assassination involving at least three assailants. The pattern of injuries indicates both planning and intense emotional involvement. Our findings provide the first genetic identification of a medieval royal, and resolve a century-old archaeological question, and illustrate the power of integrating multidisciplinary methods to confirm historical hypotheses and reconstruct violent deaths from the past with unprecedented detail.

**Teaser:** With unprecedented details, this study shows the impact of integrating multidisciplinary methods to confirm historical hypotheses and reconstruct violent deaths from the past.

## Introduction

### Historical and archaeological background

Béla (c. 1245–1272), Duke of Bosnia and Macsó, was the second son of Princess Anna (1226–c 1274), daughter of King Béla IV (reigned 1235-1270) of Hungary, and Rostislav (1219-1263) of the Rurik dynasty, Prince of Halych. Duke Béla first appeared on the political scene in 1265, when his grandfather (King Béla IV) and his mother (Princess Anna) launched a war against the younger king Stephen V. The war was led by Duke Béla of Macsó, who was around 20 years old at the time (1). His army suffered a devastating defeat, after which Béla IV made peace with the younger king Stephen V. In 1270, after the death of Béla IV, Princess Anna fled to her son-in-law, King Ottokar II of Bohemia, who then invaded Hungary. However, King Stephen V and Duke Béla repelled Ottokar and forced him to make peace. Duke Béla’s support for Stephen V stabilized the duke’s political position, but it changed after the unexpected death of King Stephen V in 1272. In opposition to the underage Ladislaus, son of King Stephen V, some of the lords supported Duke Béla as a candidate for regent or even king. Despite this, in September 1272, Ladislaus was crowned as King Ladislaus IV. To resolve the conflict, Duke Béla was invited to a council at the nunnery on the Island of Hares in November. But instead of a council meeting, an assassination plot was arranged. Duke Béla was brutally massacred by the assembled pro-Ladislaus lords led by Henrik of the House of Héder (Henrik Kőszegi). With this unpunished act, not only was a potential pretender to the throne eliminated, but the vast estates of the childless duke were also seized and later redistributed among the king’s supporters. The earthly remains of the slaughtered Béla were laid to rest by his relatives within the walls of the Dominican nunnery on the Island of Hares (now Margaret Island, Budapest) (1).

On the 6th of April 1915, archeological excavation of the Dominican nunnery on Margaret Island was initiated by K. Salkovics (2) (Supplementary Figure 1). Three graves were uncovered in the sacristy, one of which contained the visibly dismembered skeleton of a young man. Other sources describe the area between the kitchen and the so-called well-house as the site of the graves (3). In any case, it is certain that the graves were found in the 13th-century monastery building.

### Previous biological anthropological investigations

After the excavation, L. Bartucz, the anthropologist of the Department of Anthropology at Budapest University (the predecessor of the Department of Biological Anthropology at Eötvös Loránd University) investigated the remains. Based on his unpublished results, the skeleton (Figure 1, Supplementary Figure 2) belonged to a 20-25-year-old male, and 23 cut-marks were observed on the bones. Bartucz argued that the individual had not died in a duel but was likely attacked by several assailants from multiple directions; even after falling to the ground, the body appears to have been further mutilated Based on Bartucz’s theory, the remains belonged to Duke Béla of Macsó (4, 5).

**Figure 1.**
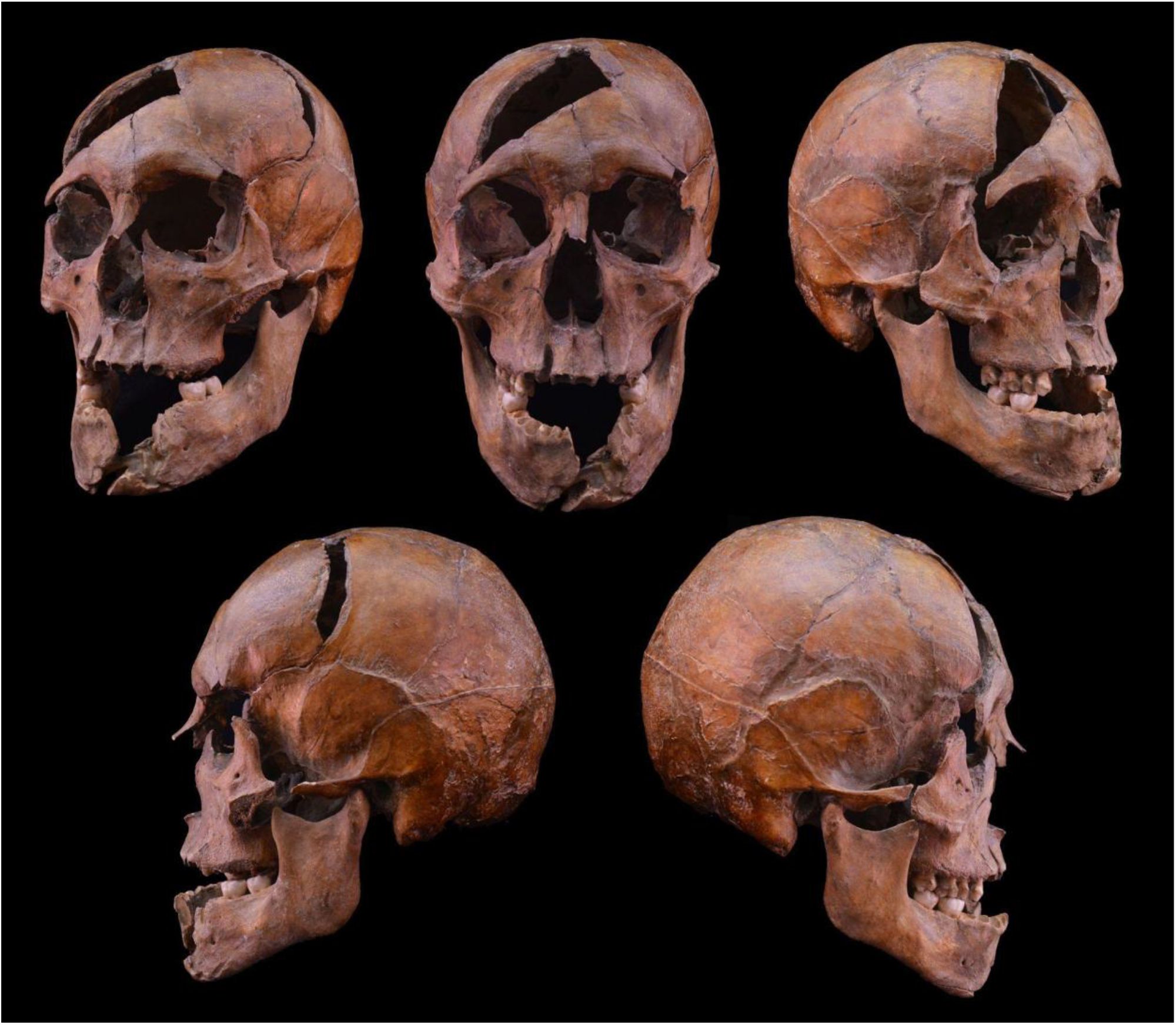
The skull of the investigated individual from the 13th century Dominican monastery on Margaret Island, Budapest.

Bernert and Buzár published the preliminary results of the investigation of the postcranial bones without the detailed investigation of the skull. They estimated the male’s age at death to be 23(±1) years. They suggested that the male was probably first stabbed in the spine from behind, which may have forced him to the ground. He was then slashed by assailants from several directions: on his hands, legs, head, and trunk.

### The aim of the study

Our study presents the results of a multidisciplinary analysis of the skeletal remains found in a Dominican nunnery on Margaret Island, aiming to reconstruct the circumstances of the individual’s life and death and establish personal identification.

## Materials and Methods

### Ethics

This research complies with all relevant ethical regulations. While archaeological skeletal remains are not considered human subjects and thus are not covered by Human Subjects regulations, we obtained formal permission for analysis of all samples from local authorities (Department of Anthropology, Hungarian Natural History Museum; Department of Biological Anthropology, Institute of Biology, Faculty of Science, Eötvös Loránd University). All samples investigated were analyzed with the goal of minimizing damage.

### Material

The skull (Figure 1) is housed in the Török Aurél Collection of Department of Biological Anthropology of the Faculty of Science at Eötvös Loránd University, while the postcranial bones (Supplementary Figure 2) are curated in the Hungarian Natural History Museum (Budapest, Hungary).

The preservation of the bones is generally good, though the lumbar spine, several phalanges, and ribs are missing. The bone surfaces exhibit minimal weathering; however, the skull interior showed extensive reconstructions from earlier treatments involving glue, paper, and plaster. The exterior mainly remained unchanged except for a possible varnish application, indicated by a visible surface shine.

### Biological Anthropology, Forensic Anthropology

Sex, age, and stature were estimated following Brůžek et al. (6), Sjøvold (7), and by the Transition analysis (TA2) (8). Craniofacial bones were scanned using a dental CT scanner (3D Accuitomo-XYZ SliceView Tomograph; J. Morita, Japan).

All bones were examined for mechanical damage, and assessments were conducted to differentiate between antemortem, perimortem, and postmortem origins. The lesions were analyzed for forensic information using macroscopic and radiologic imaging inspections. Diagnostic marks on the damaged surfaces were scrutinized to determine directions, angles, probable movement vectors, and the likely positioning of the involved parties by fitting tool proxies to the lesions. Where possible, the sequence of wound infliction was determined based on overlapping or interruptions of fractures. All injuries were also mapped onto an assembled artificial skeleton, which was used to reconstruct plausible poses and positions of the victim and the directions of the attacks (9, 10).

### Dental calculus microremain analysis

First, the teeth were visually investigated to acquire information about the diet from the dental plaque. During the sample-collecting process, powder-free gloves were worn in starch-free conditions. The series of teeth in the jaw and the spaces between the teeth were thoroughly brushed with an original hard bristle toothbrush to remove the modern contamination using ultra-pure water (UPW). The large deposits of calculus were carefully removed with a dental tool. After cleaning, the samples were processed based on King et al. (11). Retrieved calculus fragments were placed in 2 ml Eppendorf tubes, and 1 ml 10% hydrochloric acid was added to dissolve the matrix of the calculus fragments and release microremains. Stirring in acid solution lasted until no more calculus fragments were visible (15 min to 2 hours), and samples were centrifuged at 3000 rpm for 5 min. The supernatant was pipetted out, and after adding UPW, the tube was centrifuged three times.

Finally, ethanol was added, and after centrifugation and pipetting out the supernatant, the samples were dried at 30 °C. The powder-like samples were resolved by adding 20 ml of immersion oil and vortexed. The suspension was dropped on slides, and the cover slips were fixed with clear nail polish. Microremains were photographed and identified under a Nikon Eclipse polarizing light microscope at a magnification 400x. The microremains found in the dental calculus were compared with the research site’s reference microremains database (Isotope Climatology and Environmental Research Centre, Hungary). Microphotos of the samples were archived with laboratory code ML207245.

### Radiocarbon measurements

The collagen fraction of the selected bone samples, devoted to ^14^C and stable carbon and nitrogen isotope measurements, was prepared and measured at the HUN-REN Institute for Nuclear Research, Debrecen (12, 13). Prior to the AMS ^14^C analysis, samples from a rib and the occipital bone were taken. For control measurement, a sample was delivered to the University of Georgia, United States. As some influence of contamination originating from preservatives was assumed, various pretreatment protocols were suggested and tested for purification of the bones. These ranged from the modified Longin method through ultrafiltration (VIVASPIN 15R, 30,000 MWCO HY) to a complex treatment consisting of all the physical and chemical steps, supplemented with Soxhlet-extraction (33). In one case, the outer and inner layers of the bone were also separated and pretreated independently. For measurement, a small aliquot of freeze-dried gelatin obtained from each sub-sample was combusted with MnO2 reagent and the CO2 was then purified and graphitized. The bioapatite fraction of the rib was pretreated in the laboratory according to the standard protocol of cremated bones, i.e. after chemical cleaning, the structural carbonate of the inorganic part was extracted by adding 5 ml 85% orthophosphoric acid.

The AMS analysis of graphites was performed by the EnvironMICADAS AMS instrument at the INTERACT laboratory (14). As control organic samples, a blank bone (Ursus spelaeus bone with infinite ^14^C age) and internal standard bone (Bos primigenius prim. tibia from the Early Neolithic) was also applied for blank correction and checking the feasibility of the preparation process. In the case of the bioapatite measurement, international IAEA C1 (marble) and C2 (travertine) standards were applied. For the calibration and testing the possible freshwater reservoir effect (as deltaR shifts of the years) of the samples, the OxCal software (v4.4) (15) was used and the IntCal20 radiocarbon calibration curve was considered (16, 17). The calibrated date ranges are reported with 2σ probability range.

#### Stable carbon and nitrogen isotope analyses

For stable carbon and nitrogen isotope ratio analysis, repeated aliquots from the same gelatinized collagens produced by the different cleaning methods were combusted in ultraclean aluminum cups using an elemental analyzer interface (Thermo Scientific™, EA IsoLink CNSOH). Control measurements of IAEA 600 (caffeine) international, in addition, sulphanilamide laboratory standards were applied for calibration of the δ13C and δ15N values (13). The stable isotope ratios were measured using a Thermo Finnigan Delta Plus XP isotope ratio mass spectrometer. Repeated measurements were applied to achieve the precision of ±0.2‰. The stable C/N ratios of the prepared samples do not imply heavy contamination, so the stable isotope results (Supplementary Table 1) can be used in subsequent analyses.

#### Strontium isotope analyses

The enamel of two left lower molars (M1, M2) has been sampled in three points of the tooth surface: at the crown cusp, at the middle of the crown and at the enamel-root junction. Strontium (Sr) sample solutions for isotope ratios were measured by a Neptune Plus MC-ICP-MS (Thermo Scientific) equipped with an Aridus-3 (CETAC) desolvating nebuliser system (dry plasma) at ICER) (18). The raw measurement data were normalized to a standard solution of 35 ppb prepared from the NIST SRM987 international Sr standard (Nat. Inst. of Stand. and Techn.). The ^87^Sr/^86^Sr ratio was corrected for instrumental mass discrimination using ^88^Sr/^86^Sr = 8.375209, as well as by applying an interference correction for ^87^Rb+ and ^86^Kr+ with ^85^Rb+ and ^83^Kr+, respectively. All values were normalized to the reported value of 0.710248 for NIST SRM 987 (18).

### Genetic testing

The laboratory work for genetic testing was conducted at the Institute of Archaeogenomics, Research Centre for the Humanities, ELTE (Budapest, Hungary), and the Department of Genetics, Harvard Medical School. All operations for the uniparental investigations were conducted within the dedicated ancient DNA laboratory of the Institute of Archaeogenomics, Research Centre for the Humanities, ELTE in Budapest, Hungary. The whole genome capture and sequencing was undertaken at the Laboratory of the Harvard Medical School, after the sample preparation was carried out at the Department of Evolutionary Anthropology of the University of Vienna.

#### Laboratory work in Budapest for uniparental analysis

The sample BPDOM01 was drilled from the left petrous part of the temporal bone from the basis of the intact skull in Budapest.

DNA was extracted from petrous bone powder using established protocols in Budapest, incorporating laboratory-specific modifications. DNA extraction followed the procedure outlined by Dabney et al. (19), with minor adjustments as highlighted by Lipson et al. (20) from 50 mg bone powder.

##### Mitochondrial DNA workflow

DNA library preparation involving partial (half) uracil-DNA-glycosylase treatment followed the protocol outlined in Rohland et al. (21), with the cleanup step optimized for robotic automation. Partially double-stranded and barcoded P5 and P7 adapters were employed. Amplification of the libraries was conducted using TwistAMP (TwistDX). The resulting amplification products were purified using AMPure Beads Purification (Agilent). For capturing target sequences encompassing the entire mitochondrial genome an in-solution hybridization method was adopted, as described by Csáky et al. (22). The captured samples were indexed using universal iP5 and unique iP7 indexes. Next-generation sequencing was conducted utilizing the Illumina MiSeq System (Illumina) employing V3 (2 × 75 cycles) sequencing kits.

##### Y STR workflow

We investigated short tandem repeats (STRs) on the Y-chromosome using the AmpFlSTR® Yfiler® PCR Amplification Kit (Applied Biosystems). Amplification followed the manufacturer’s protocol with a single modification: the PCR was run for 34 cycles. This modification was required because the specimen is of ancient origin, characterized by very low DNA concentration in the extract and a predominance of highly fragmented template DNA. Amplicons were fluorescently labeled with 6-FAM™, VIC®, NED™, and PET®, and were sized against the GeneScan™ 500 LIZ® Size Standard. An allelic ladder was included in every run. We performed three independent PCR reactions on the sample and three independent capillary electrophoresis and data acquisition.

PCR products were analyzed on a 16-capillary ABI 3130xl Genetic Analyzer using POP-7 polymer, 50 cm capillaries, and the G5 filter set, with a 35-500 bp sizing range. All three analyses were performed under identical instrument and run conditions, no parameter changes between runs.

#### Laboratory work at Harvard for whole genome analysis

Bone powder was prepared from cochlea at Pinhasi Lab, Vienna following the Pinhasi et al. (23) method. DNA was extracted from 35 mg of powder using an automated (‘robotic’) procedure using silica magnetic beads and Dabney Binding Buffer on the Agilent Bravo NGS workstation (24). USER-treated single-stranded library was produced using automation and then enriched in-solution for sequences overlapping 1.24 million SNPs. (25, 26). The sequencing platform for 1240k capture data was Illumina HiSeq X10, 2x101 bp read length.

### Bioinformatic data analysis

#### Mitochondrial data analysis

We processed the raw mitogenome sequence data using the same in-house pipeline outlined in Gerber et al. (27), utilizing the PAPline package with default settings (28). Contamination levels of the mtDNA were estimated using the ContamMix software (26), for results, see Supplementary Table 7. The mtDNA haplogroup was determined by Haplogrep (29, 30). Comparing to the rCRS sequence, we excluded the following ambiguous base positions and heteroplasmic hotspots: 42, 57, 291–317, 447–458, 511–524, 568–573, 594–597, 1718, 2217–2226, 3106–3110, 3159–3167, 5890–5894, 8272–8281, 16,184–16,193. For mitochondrial phylogenetic analysis, method described in Csáky et al. (22) was followed, using all known and available sequences assigned to U3 haplogroup, on the figure keeping only the U3b3 sub branch (Supplementary Table 3).

#### Y chromosomal data analysis

The Y STR data analysis was carried out with GeneMapper® ID Software v3.2.1 (Applied Biosystems). Allele-calling threshold was 150 RFU. To examine the STR variation within the Y-chromosomal haplogroup, Median Joining (MJ) network was constructed using the Network 10.2.0.0 program and the figure was drawn with the Network Publisher 2.1.2.5 program.The DYS385 locus was excluded because the Network program cannot handle the duplicated loci (DYS385a, DYS385b).

Yleaf (31) was applied for Y-chromosomal haplogroup determination from next generation sequencing data (Supplementary Table 2).

#### Whole genome data analysis

##### Raw data processing

The sample underwent sequencing in Harvard Laboratory to produce raw paired-end reads. Paired-end reads were merged into single-molecule sequences based on overlapping regions. Single-end reads were aligned to the hg19 human reference genome (32) and the Reconstructed Sapiens Reference Sequence (RSRS) mitochondrial genome using the *samse* aligner of BWA. Duplicate molecules were identified and marked based on barcoding bins, start/stop positions, and orientation. The computational workflows, along with the specific parameters used, are publicly available on GitHub: ADNA-Tools and adna-workflow (33).

For variant calling, a pseudo-haploid approach was applied to targeted SNPs, wherein a single base was randomly selected from the possible alleles at each position. Filtering criteria included a minimum mapping quality of 10 and a base quality of 20, with additional trimming of two base pairs from both the 5’ and 3’ ends of reads to mitigate damage-related artifacts.

##### Contamination prediction

The ratio of male X-chromosomal contamination was measured, with both ANGSD (version 0.939-10-g21ed01c, htslib 1.14-9-ge769401) (34) and hapCon (hapROH package version 0.60) (35). All parameters used in both methods were the default settings recommended in the official documentation. The starting file for the former method was a samtools mpileup (version 1.10, htslib 1.10.2-3ubuntu0.1) output with mapping quality 25 and base quality 30. ANGSD doCounts was run on the X-chromosomal region from position 5500000 to 154000000, with mapping and base qualities 25 and 30 respectively.

##### Imputation and IBD analysis

Imputation was done with the GLIMPSE1 software suite (36). During the process, we mainly followed the developers’ official GRCh37 tutorial (37) and used the v1.1.1 versions of the precompiled static binaries distributed on github (38).

The reference panel used was the 1000 Genomes Project (30X from NYGC) on hg19/GRcH37. Genotype likelihoods of the samples were computed with a standard method involving bcftools mpileup and bcftools call (version 1.10.2). Base- and mapping quality (flags -Q and -q) were both set to 20. As a difference from the official workflow, chromosomal level imputation regions were determined on a per sample basis, with the GLIMPSE_chunk_static executable, using default parameters (--window-size 2000000 -- buffer-size 200000).

To identify identity by descent segments among samples we used ancIBD (version 0.5) (39) with the same process and parameters described in the software’s official documentation (40).

##### Clinical and pigmentation variant calling

Nuclear variants with clinical significance and pigmentation variants were analysed with the variant calling tool from the PAPline package (27). The panel contains 18738 variants, of which 61 are pigmentation markers, and the rest are clinically relevant positions.

## Results

### Radiocarbon investigation

The collagen fraction of bone samples was prepared in multiple ways (n=7), yielding conventional ^14^C dates ranging from BP 874±19 to BP 922±17, corresponding to a summarized calibrated age range of approximately cal AD 1030-1230 at a 95.4% probability level (Supplementary Figure 3). We could not observe significant differences between the results obtained by the different pretreatment methods for the rib or skull fragments. Thus, a freshwater reservoir effect test, related to aquatic-derived food, was performed applying an age shifting of 50-150 yrs towards the younger calendar dates. The structural carbonate of the bioapatite fraction of a rib was also prepared and measured, resulting in a conventional age of BP 841±18 (AD 1170-1260 at 95.4% probability level).

### Biological Anthropology

The transition analysis aging method integrates findings from 24 skeletal traits. The estimated age-at-death ranges between 19.8 and 30.9 years (95% confidence, corrected data), with a maximum likelihood estimate of 24.3 years. The determined sex (male) based on morphological traits of the cranium (Figure 1) as well as the metric characteristics of the pelvic bone, corresponds with the genetic sex. The estimated stature of the individual was 178.8 cm, based on the average femoral length.

Slight marginal osteophytes can be seen on both condyles of the left femur and right tibia. Slight hypertrophy of the subchondral bone (5.7 x 3.9 mm) can be seen on the medial side of the right patellar articular surface. Slight marginal bone “lipping” is visible on the upper articular processes of the 4th-9th thoracic vertebrae. The insertion sites of the muscles and tendons are slightly expressed on the tuberosity of the right radius, on the deltoid tuberosity of both humeri, on the linea aspera of left femur, and on the soleal line of both tibiae. The lateral and anterior surface of the vertebral bodies are hypervascularized.

Healed mild periosteal lesions appeared on the medial surface of the mid-third part of both tibiae, and on the mid-third part of the left fibula.

### Odontology

Eight teeth are intact and properly positioned in the mandible, surrounded by healthy periodontal bone. Upper left 1^st^ premolar and 1^st^-2^nd^ molars are fractured in the region of the tooth neck, presumably due to forced impact immediately before death. The plane of fracture of the two molars is the same as the plane of the cut of the zygomatic bone, the zygomatic process of the maxilla and the vestibular bony process of the upper jaw around those molars (Supplementary Figure 4A-B). A non-dislocated fracture of the mesiobuccal root of the left upper first molar tooth was also observed (Supplementary Figure 4C).

### Dental calculus analysis

Presumably, due to the individual’s young age, the amount of calculus was not particularly high (14.3 mg). 791 microphotographs were taken, showing more than one thousand microfossils (41). It was estimated that 80% of the microfossil remains were microcharcoal, starch and miscellaneous unidentified plant and animal remains, with a smaller portion of fibers, phytoliths, pollen, animal and plant egg cells and spores, and fungal remains (Supplementary Figure 5). The size and shape of the starch granules predominantly suggest they came from the endosperm tissue of grains belonging to the Triticeae tribe, namely wheat (*Triticum*) and barley (*Hordeum*). In addition, several long plant hairs (trichomes) were detected that are characteristic of the wheat grains (Supplementary Figure 6). The starch grains’ identifiability was limited because they bore marks of grinding, cooking, and baking (42). Regarding the size, morphological and light refraction characters of the starch grains and the material of the gelatinized starch in the dental calculus were similar to the reference material of the boiled wheat grits and the baked wheat bread (Supplementary Figure 6). Yeast cells may also have integrated into the calculus, but it cannot be established with certainty. In the sample of the individual’s dental calculus, 17 pollen grains were detected, which is a large number (43), and numerous calcium oxalate crystals were found in the calculus matrix.

### Forensic traumatological assessment

The assessment of the skeleton reveals 26 perimortem lesions (Figure 1-2) – nine on the skull (Figure 1-3, Supplementary Figures 7 and 8) and 17 on postcranial bones (Figure 2 and 4, Supplementary Figure 9-17) – all sustained in a single violent event (Supplementary Information 1). All wounds are clearly the result of intent and interpersonal aggression. Their characteristics suggest a coordinated assault involving probably three assailants.

**Figure 2.**
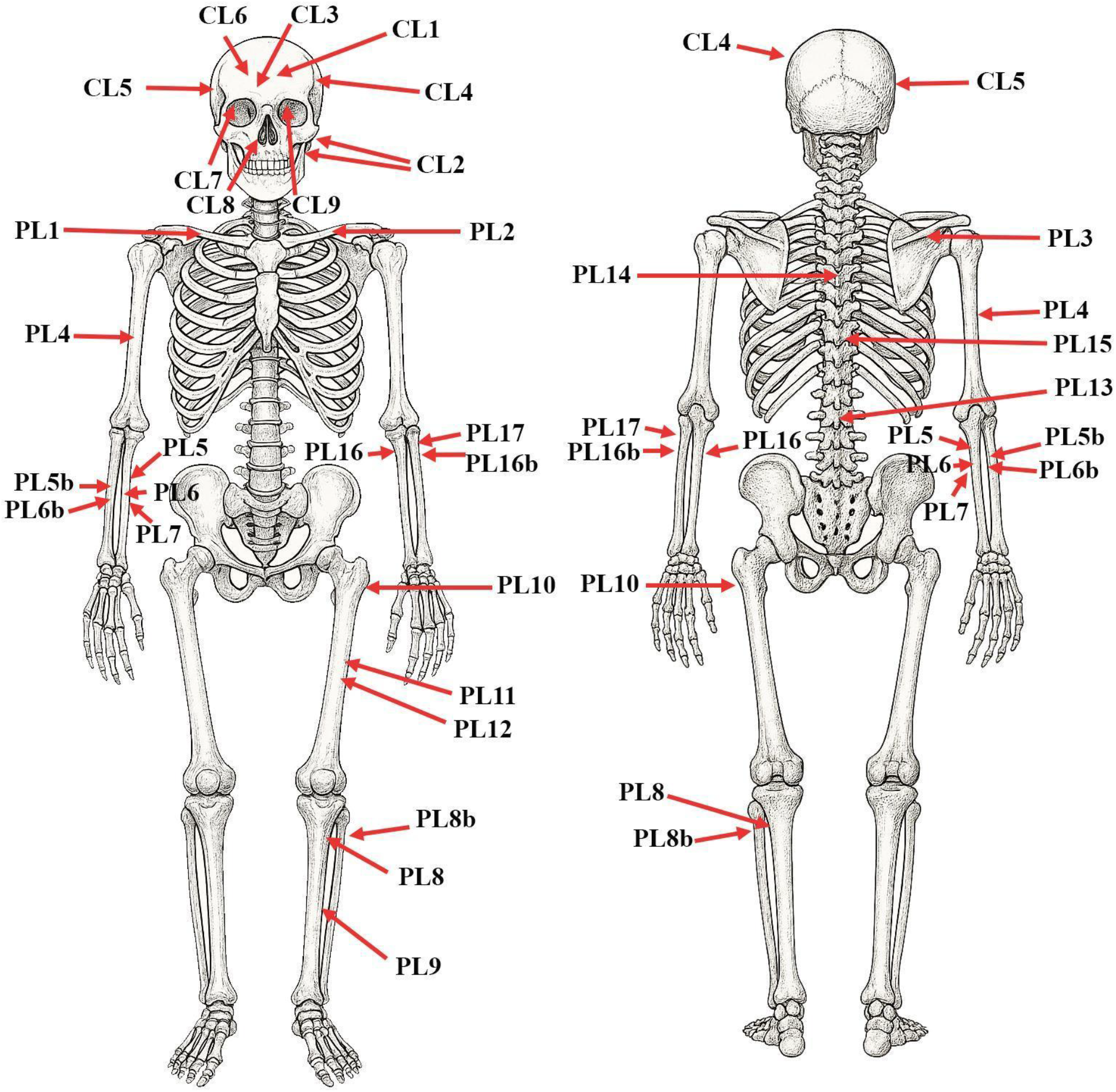
The observed perimortem lesions on the human remains. The description of the traumatic alterations can be found in Supplementary information 1.1 and 1.2. CL=cranial lesion, PL= Postcranial lesion. The drawing of the skeleton was generated using OpenAI’s image generation tools (DALL·E) via ChatGPT.

**Figure 3.**
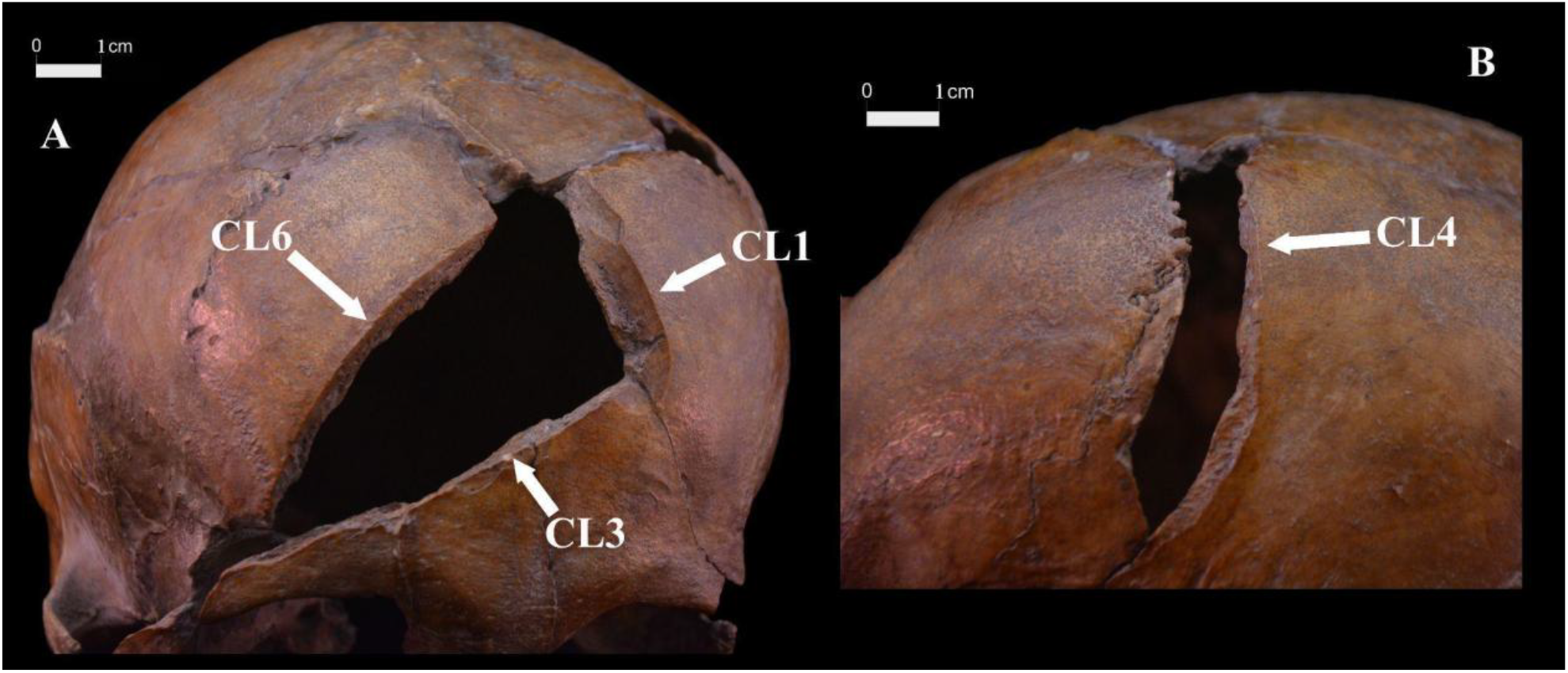
Perimortem lesions on the frontal (A) and left parietal (B) bones caused by sharp weapons. White arrows show the sharp edges of the traumatic alterations without healing. The description of the traumatic alterations can be found in Supplementary information 1.1. CL=Cranial Lesion

**Figure 4.**
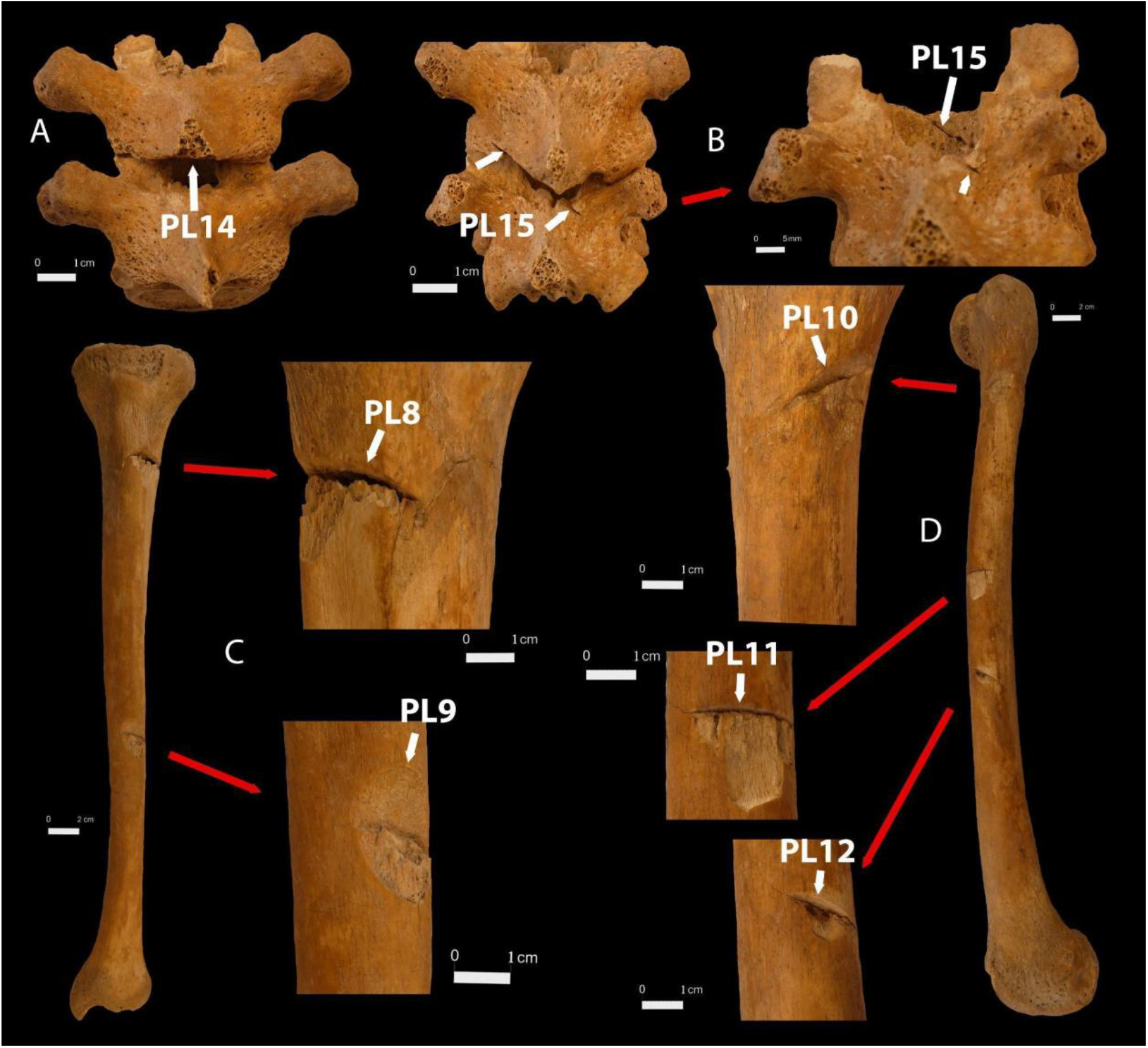
Perimortem cut-marks on the postcranial bones. White arrows show the sharp edges of the traumatic alterations without signs of healing. A: Perimortem cut marks on the spinous process of the 7th thoracic vertebra (PL14). B: Perimortem cut marks on the 10-11th thoracic vertebrae. The tip of the tool went from behind through the vertebral canal and was stopped by the vertebral body of the 11th thoracic vertebra (PL15). C: Perimortem cut marks on the anterolateral surface of the left tibia (PL8, PL9). D: Perimortem cut marks on the anterolateral surface of the left femur (PL10, PL11, PL12). The description of the traumatic alterations can be found in Supplementary information 1.2. PL=Postcranial Lesion

One attacker approached from the front; the others were flanking the victim from the left and right. The injuries, primarily on the front of the body, imply the victim faced his attackers in an open confrontation. The presence of multiple defensive wounds on the forearms confirms he was aware of the attack and attempted to defend himself.

The weapons used can be identified as two different types of blades, probably saber and longsword (the detailed arguments are shown in Supplementary Information 2). The clean and deep cut marks, especially on the skull, suggest the victim was unarmored, as body armor or a helmet would have prevented such injuries.

The reconstruction of the assault sequence (Supplementary Information 3) indicates the attack began with saber strikes to the head and upper body, followed by defensive injuries as the victim attempted to block further blows. The attackers eventually incapacitated the victim with further strikes from the flanks, continuing the assault as he fell to the ground, and delivering fatal injuries to the head and face.

The intensity of the attack clearly indicates an intent to kill, while the numerous blows to the face suggest intense emotional involvement, such as rage or hate (“overkill”). However, the coordinated nature of the attack implies premeditation, suggesting this was at least partly a planned assassination.

### Strontium, stable carbon and nitrogen isotope ratio

Based on the measurement results of the Sr isotope ratios the 1^st^ molar forms a group (mean: 0.710363±0·000184), the 2^nd^ molar forms a different group (0.709769±0·000111) (Supplementary Figure 18). Strontium isotope values of the first molar show that Béla was born and spent his early childhood in a different geographic place, as where he was buried, characterized by more radiogenic values, such as the regions of Vukovar or Sirmia (44), which were part of the Banate of Macsó. However, the exact and certain localisation of the territory where the investigated male was born and spent his early childhood, is not possible because of methodological limitations. In the Carpathian Basin and surrounding territories, more regions have the same or very similar strontium isotope values (44).

During late childhood, the subject moved to a different region, possibly in the hinterland of Budapest. The ^87^Sr/^86^Sr value measured on the rib appears compatible with the local baseline, although bone tends to undergo a diagenetic alteration and contamination with the depositional soil.

The d13C of -19.3‰implies a primarily C3-type plant-based diet. The stable nitrogen isotopic result (d15N) of 11·8‰is situated above the range 6-10‰, which is characteristic for humans consuming largely terrestrial plants and animals (45). It suggests that animal-, and possibly aquatic-derived protein might have made a notable contribution to the diet of this individual.

### aDNA analysis

The maternal and paternal relatives of Béla of Macsó, are well-known and documented. There are published genetic studies available from both the paternal Rurik side and the maternally related Árpád Dynasties. These allowed us to make comparisons at the whole-genome level and for Y-chromosomal and mitochondrial uniparental lineage.

#### Analysis of the paternal lineage

For our analyses of the Y-chromosomal paternal side, we included modern samples from the *Rurikid Family Tree DNA Projec*t (46) representing descendants of the Rurik Dynasty, as well as the whole-genome data of Prince Dmitry Alexandrovich (c. 1250–1294) (47), who was a Grand Prince of Vladimir (northeastern principality of the Rus) and an eighth-degree relative and contemporary of Béla of Macsó from the Rurik dynasty. Prince Dmitry Alexandrovich’s great-grandfather (Vsevolod III, Rurik dynasty) had a son (Sviatoslav III, Rurik dynasty) who was the great-great grandfather of Béla of Macsó. Firstly, we investigated the short tandem repeats (STRs) of the Y-chromosome. Three independent reactions resulted in an STR profile based on 13 Y-STR loci, which suggested that DNA sample BPDOM01 belonged to the N1a1-M46 haplogroup (*nevgen*.*org*). Further downstream subgroups can be anticipated from the STR matches detected with the Rurikids and based on Y-chromosomal SNPs from the whole-genome analysis, BPDOM01 most probably belonged to the subgroup N1a1a1a1a1a1a (N1a-L550, ISOGG 2020 Version: 15.73 (Supplementary Table 2). Based on publicly available Y chromosome STR data with the participants of the Rurikids Dynasty project, we selected those families (branches) that had members who shared the most STR alleles with the sample BPDOM01. A median-joining network analysis with these Rurikid branches was performed with Network 10.1.0.0 (Figure 5A). According to the database of the FamilyTreeDNAa’s Rurikid Dynasty DNA Project (46), the 13 Y-STR profile of BPDOM01 has 24 exact matches among the tested descendants of the Rurikids and Varangian families. Furthermore, it has a perfect match with the anticipated Rurik prince’s Y-STR profile (lived ca. 830–879 AD), who was a paternal ancestor of Béla of Macsó in 12 generations distance. Among the present-day descendants, members of the Rurikids, Rurikid Princes, Varangians - ‘Rurikids closest cousins’ and ‘Proto-Rurikids’, and Finno-Karelian Branches share all 13 STR from the typed 13 markers of BPDOM01. The sub haplogroups of these individuals belong to Y10931 (N1a1a1a1a1a1a7a3a∼) and Y4339 (N1a1a1a1a1a1a7a∼) both located downstream from L550 on the Y tree (Supplementary Figure 4). This has been confirmed by the Yleaf analysis of genomic data, the S431 derived marker defining the N-L550 haplogroup is covered by three reads. This haplogroup classification supports the theory that the BPDOM01 sample belongs to a descendant of the Rurik Dynasty. The strongest evidence of his Rurikid origin is provided by the whole-genome-based Y-chromosomal haplogroup classification of Prince Dmitry Alexandrovich, which is N1a1a1a1a1a1a7a3b∼ (N-VL11 terminal SNP), consistent with our results.

**Figure 5.**
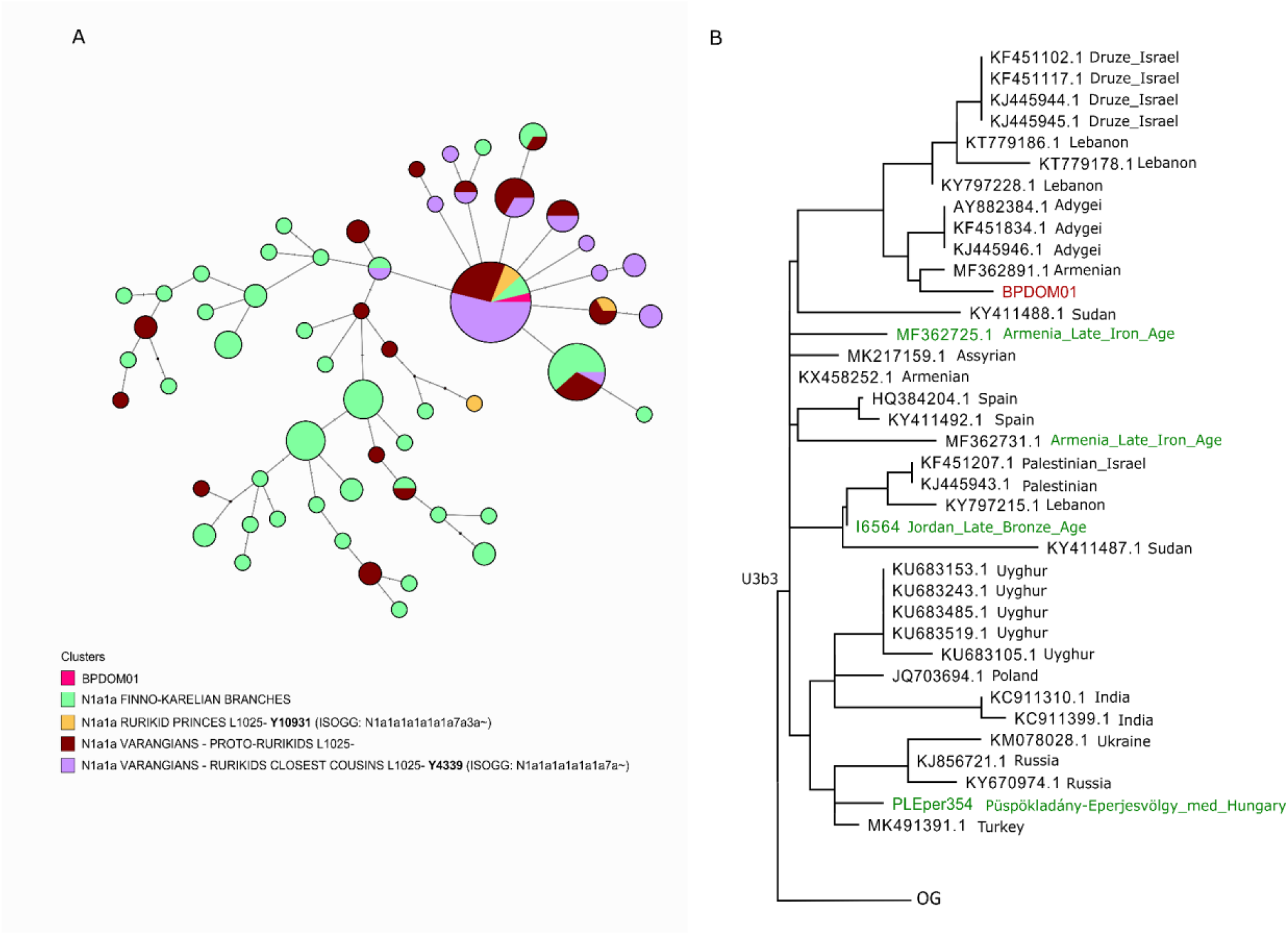
Phylogenetic networks of the uniparental lineages of sample BPDOM01. Part A: Median Joining Network based on 11 Y-STR data of the closest N1a-L550 individuals’ clusters from the Rurikids project. The closest individuals to BPDOM01 on the Median Joining Y network are derived for the SNP L550, but ancestral for the L1025 marker. Most of them are located under the VL15 and Y10931 “Rurikid” branches on the N phylogenetic tree, downstream from L550. Part B: Part of the U3 mitochondrial phylogenetic tree, showing the entire U3b3 branch. On the U3b3 tree, the most closely related lineages to the BPDOM01 sample are from present-day Armenia and the north-western Caucasus.

#### Analysis of the maternal line

The maternal line of BPDOM01 can be classified as the mitochondrial sub-haplogroup U3b3, with an average mitogenome coverage of 66x. No identical mtDNA haplotype to BPDOM01’s lineage was detected in published datasets. The U3b3 haplogroup has its present-day highest frequency around the Black Sea. It appears in ca. 3-6% in Turkey, populations of the Caucasus and Iraq, Iran today (48). This mtDNA subgroup’s earliest appearance in ancient databases is in Minoan Greece, Crete (1871 BC), the Late Bronze Age Levant (1276 BC; today’s Jordan) and the Iron Age Armenia (526-494 BC) (49, 50). Further ancient samples with the same maternal sub haplogroup are known from the following regions and periods: Hungary (Püspökladány-Eperjesvölgy, 975-1075 AD), Poland (Piast dynasty, 1040 AD), Turkey (Byzantine culture, 1163 AD). The phylogenetic tree of all available U3b3 mitogenomes is shown in Figure 5B, with the closest matches being modern Armenian, Adygei, Lebanese and Druze sequences (Supplementary Table 3). The maternal grandmother of Béla was Maria Laskarina (c. 1206-1270), a daughter of the Byzantine Emperor Theodor I., and Anna Komnene Angelina (c. 1176-1212), who was the Empress of Nicaea and daughter of the Byzantine Empress Euphrosyne Doukaina Kamaterina (c. 1155-1211). The U3b3 lineage aligns with the here presented maternal genealogy leading to Euphrosyne Doukaina as a rare but widespread lineage from the Caucasus to the Near East, likely prevalent in the Early Medieval Anatolia and the Aegean – South-Pontic world, from which Béla’s maternal ancestors came.

#### Autosomal DNA analyses

We present the genome-wide single nucleotide polymorphism (SNP) data (545,606 SNP covered) from 1240k SNP capture of BPDOM01 and comparative samples in a principal component analysis (PCA) (Figure 6A, Supplementary Table 4). For the PCA analysis, ancient data were projected on the modern European genetic variation using the Human Origin panel as a reference dataset (included in the AADR database) (51). BPDOM01 aligned closest to modern Greek, Turkish and Moldavian samples in the PC1-PC2 (principal components 1 and 2) space, slightly extending towards the southeastern populations, which are representative of the present-day Caucasian groups. The closest samples originate from the medieval era of today’s Hungary and Romania and medieval/ early modern Italy.

**Figure 6.**
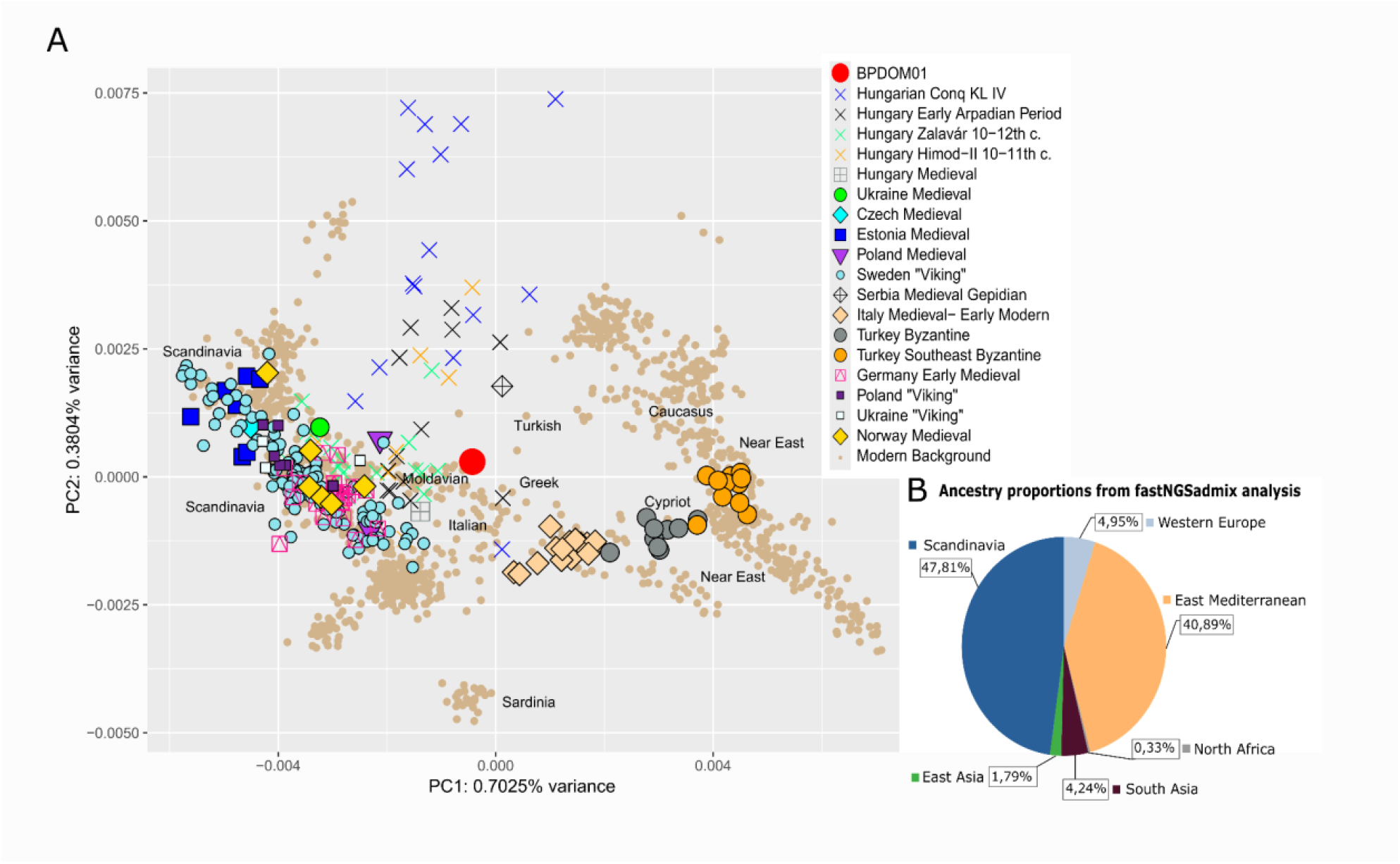
European PCA and supervised genomic ancestry proportions of BPDOM01. A: PCA results with genotypes from medieval European populations. 560k SNPs were used for the calculations, performed in the smartpca program. The scatterplot was generated in R. B: fastNGSadmix analysis with an ancient reference panel following the concept presented at Vyas et al. (53)

To explore Béla of Macsó’s genomic composition, we applied the fastNGSadmix software. The fastNGSadmix program is an efficient and reliable tool for determining admixture proportions from NGS data of a single individual. It utilizes genotype likelihoods for its analyses, comparing the input sample to a set of predefined reference populations. This method maintains high accuracy even at low sequencing depths and compensates for biases introduced by small reference populations (52). Similar to those presented in Vyas et al. (53), we used a custom panel of ancient 4th–8th-century reference individuals. This panel was constructed with a similar geographic distribution to the 1000G populations to form seven ancient panel groups, such as Western Europe, Sub-Saharan, East Mediterranean, North Africa, South Asia, East Asia, and Scandinavia as in the premade reference panel of the fastNGSadmix program. The results align with the presumed origin of Béla of Macsó, having Byzantine (Eastern Mediterranean) ancestry on the mother’s side and Rurik (Scandinavian) ancestry on the father’s side (Supplementary Figure 6B) (52).

Applying ancIBD, the identity-by-descent (IBD) tool of Ringbauer et al. (39), the genomes of Dmitry Alexandrovich and BPDOM01 share a total of 118 cM DNA fragments with the largest matching chunk over 43 cM (Supplementary Table 5, Figure 7A and B). This result confirms them to be at least eighth-degree relatives, consistent with the historically recorded kinship between Dmitry and Béla of Macsó.

**Figure 7.**
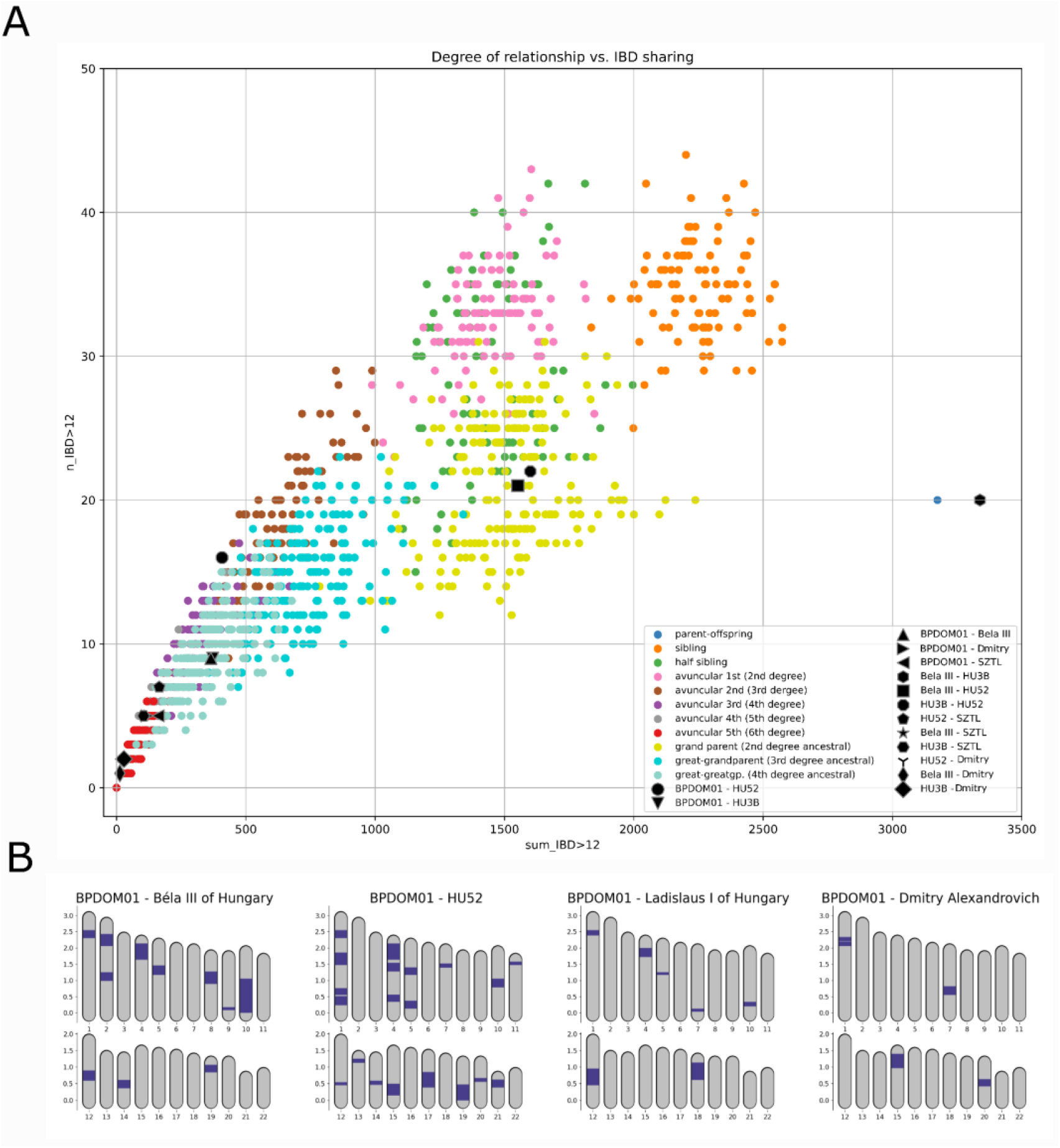
Degree of relationship vs IBD sharing and IBD sharing with known medieval personals from Hungary and Russia. Fig. 5A: Degree of relationship on simulated data with IBD sharing segments. Fig. 5B: Karyotype Plot of IBD sharing segments. This function of ancIBD depicts IBD between a pair of individuals along their chromosomes. The simulated data for comparison of relatedness was gained from Ringbauer et al. (39)

Besides the sample of Prince Dmitry Alexandrovich, we incorporated into our ancIBD analysis the whole-genome sequences of Béla of Macsó’s maternal great-great-grandfather, King Béla III (1148-1196) of Hungary from the Árpád dynasty, further samples from the Coronation Basilica in Székesfehérvár, and a sample from the herma (skull relic) of St. Ladislaus from the Árpád dynasty (c. 1040-1095) (54–56).

The ancIBD analysis (Figure 7A and B) reveals that the examined BPDOM01 sample is a fourth-degree relative of King Béla III, and approximately twice as distant from St.

Ladislaus (55, 56), which supports the hypothesis that he was Duke Béla of Macsó (Supplementary Table 5). In the ancIBD run, we verified the identity of the data labelled under the codes BelaIII and HU3B. The former is from the publication by Wang et al. (55), and the latter is from the publication by Varga et al. (56). These are two different samples, both originating from Béla III (see Supplementary Table 5).

The ancIBD showed that the BPDOM01 sample is probably a fourth-degree relative on an avuncular line of the HU52 sample from the Árpád dynasty, found in the Coronation Basilica in Székesfehérvár. Considering the historical and genetic data, we conclude that the skeleton from grave III HU52 may have been identical to Prince Andrew of Halych, the son of King Andrew II. Following the rationale and terminology employed by Olasz et al. (54), we refer to the sample included in the present analysis as HU52, originating from Grave III of the Coronation Basilica in Székesfehérvár, given that the published sequence used in our study (56) is also associated with this identifier. It is important to note, however, that this individual is not identical to the one labelled II/52 in the original excavation records of the basilica, who was identified as a fetal skeleton.

We applied a clinical variant calling and pigmentation analysis on sample BPDOM01 with the variant calling tool from PAPline (27). Through the 1240k SNP capture, altogether 32 pigmentation markers were captured and detected with a 3.7x average coverage. Overall darker pigmentation and darker skin, no melanoma risk or sunburn sensitivity characterized BPDOM01 and freckles were not characteristic, possibly having brown curlier hair and light brown eyes. No disease-associated genotypes were identified through the examination of 925 clinical variants (see Supplementary Table 6).

## Discussion

The radiocarbon results of bone collagen indicate an earlier date than the recorded murder of Duke Béla. However, the building in which the skeleton was found is part of the Dominican monastery on Margaret Island, which was founded only in 1259 AD. As the first buildings of the monastery were built between 1246-1255, the burial could not have taken place earlier than 1246. Our ^14^C measurements were conducted seven times on the collagen fraction of the bones, and verified by independent laboratories, and the results are consistent. As the younger age of the bioapatite fraction suggests, the discrepancy between the radiocarbon results and historical data can likely be explained by the effect of the individual’s diet. Moreover, the higher d15N value also supports the possibility of reservoir effect shifting the real ^14^C age toward an older date (57, 58). Although the dental calculus microremain analysis results did not provide evidence of this type of diet, it cannot be ruled out that the level of ancient carbon in the diet was sufficiently high to influence the radiocarbon results, causing a reservoir effect of 100-200 years for the collagen fraction and 50-100 years for the structural carbonate of bioapatite. Since our genetic analysis confirms that the investigated male belonged to the highest level of society (as a member of the Árpádian Dynasty), access to expensive (luxury) and peculiar foods (e.g. seafood, shells and fish), even in the middle of the Carpathian Basin is probable. Despite the radiocarbon results, our multidisciplinary data thus support the early 20th century hypothesis of L. Bartucz that the investigated remains represent Béla, Duke of Macsó (4).

The IBD analysis verified that the individual in question is a fourth-degree descendant of King Béla III of Hungary from the Árpád dynasty, supporting the hypothesis that he was Duke Béla of Macsó. His ancestry, characterized by a mixture of medieval Scandinavian, Eastern Mediterranean, and Árpád-era Hungarian components, along with his Y-chromosomal lineages matching that of the Rurikid house, validates the hypothesis derived from historical data through whole-genome analyses.

The estimated age-at-death and osteological sex of the examined individual also correspond to expectations of Béla, Duke of Macsó, who was likely born c. 1245 and died in 1272.

The stable isotope results indicate that Béla of Macsó’s diet was rich in C3 cereals (such as wheat and barley) and animal proteins, reflecting high social status. The dental calculus analysis further supports the consumption of wheat and barley, which fits into the known dietary patterns of medieval populations in the region (Supplementary Information 4) (59). The forensic traumatological investigation allowed us to reconstruct the circumstances of Duke Béla of Macsó’s death. All his wounds are clearly caused by interpersonal aggression and a coordinated assault involving probably three assailants. The brutality of the attack, the encircling of the victim with strikes from three sides, and the character of blows show the intent to leave no chance of survival for the duke. The numerous poorly aimed attacks on the victim and the degree of damage to his face suggest a strong emotional involvement of hate and rage. Thus, while possibly premeditated and planned, the murder was not committed in cold blood.

## Acknowledgments

We thank the National Centre for Biodiversity and Gene Conservation (Hungary) and the Research Institute of Karcag (Hungary) for plant reference material.

## Funding

The study was supported by the grants of the Hungarian Research, Development and Innovation Office FK142894 (ZSLSZ); the Bolyai Scholarship of the Hungarian Academy of Sciences (TH – project id: BO-783-22-8, IM – project id: 710-23-10, ZSLSZ: project id: 357-23-10). Project No. KDP-2023-C2284509 has been implemented with the support provided by the Ministry of Culture and Innovation of Hungary from the National Research, Development and Innovation Fund, financed under the 2023-2.1.2-KDP-2023-00002 funding scheme (TH). IM’ work was supported by the European Union and the State of Hungary, co-financed by the European Regional Development Fund (project of GINOP-2.3.4-15-2020-00007 “INTERACT”). The study was also supported by the “MALTHUS” project, funded by the Italian Ministry of University and Research (PRIN2022: 2022ELZECR – (CC)).

This research was supported by the Hungarian Natural History Museum and the House of Árpád Programme (2018–2023) Scientific Subproject: V.1. Anthropological-Genetic portrayal of Hungarians in the Árpádian Age (TH, TSZ, ZSB, ÁB).

Research work of NB was supported by the EKÖP-24-3-II university excellence scholarship program of the Ministry for Culture and Innovation from the source of the National Research, Development and Innovation Fund. ID: EKÖP-24-3-II-ELTE-474

## Competing interests

The authors declare that they have no competing interests.

## Data and materials availability

The results of the biological anthropological, radiocarbon, stable isotope, dental calculus microremain analysis are available in the main text and in the supplementary materials. The ancient genome sequence of the BPDOM01 sample has been uploaded to ENA (European Nucleotide Archive), under the accession number PRJEB92396.

Data required to generate all figures in the manuscript are available in the main text and in Supplementary materials.

## Notes

### Competing Interest Statement

The authors have declared no competing interest.

### Summary of Updates

We substantially revised the manuscript's structure and, based on reviewer feedback, clarified and expanded methodological details, and updated the supplementaries.

