## Supplementary Information for "Murder in cold blood? Forensic and bioarchaeological identification of the skeletal remains of Béla, Duke of Macsó (c. 1245–1272)"

**This PDF file includes:**

Supplementary Text
Supplementary Figures S1 to S18
References (1 to 23)

**Other Supplementary Materials for this manuscript include the following:**

Supplementary Tables S1 to S7

#### Table of Contents


Supplementary Information 1. Forensic traumatology 3
    Supplementary Information 1.1. Cranial lesions 4
    Supplementary Information 1.2. Postcranial lesions 8
Supplementary Information 2. Weapon characteristics 14
Supplementary Information 3. Forensic evaluation and situational reconstruction 15
    Supplementary Information 3.1. Forensic traumatological assessment 15
    Supplementary Information 3.2. Reconstruction of the situation 17
Supplementary Information 4. Diet of the Medieval Hungarians 18
Supplementary Figures 19
Supplementary Figure Legends 20
Supplementary Figure 1 23
Supplementary Figure 2 24
Supplementary Figure 3 25
Supplementary Figure 4 26
Supplementary Figure 5 27
Supplementary Figure 6 28
Supplementary Figure 7 29
Supplementary Figure 8 30
Supplementary Figure 9 31
Supplementary Figure 10 32
Supplementary Figure 11 33
Supplementary Figure 12 34
Supplementary Figure 13 35
Supplementary Figure 14 36
Supplementary Figure 15 37
Supplementary Figure 16 38
Supplementary Figure 17 39
Supplementary Figure 18 40
Supplementary Tables (Supplementary Tables are in a separate xls file) 41
Supplementary Table Legends 42
References cited in Supplementary Information 1-4 and in the Supplementary Tables 43

#### Supplementary Information 1. Forensic traumatology

With the forensic investigations for traumatology, we seek to answer the following specific questions:

- Were the previously observed lesions on the skeletal remains caused by interpersonal violence?
- If so, how many assailants were involved?
- Which implements were used to cause the observed lesions?
- Can the situational reconstruction differentiate between an open confrontation and an ambush?
- Can the evidence discern between a premeditated murder and a manslaughter driven by emotional states?

*Supplementary Information 1.1. Cranial Lesions*

(in order of probable infliction sequence)

Lesion 1 (CL1):

Description: A superficial, tangential, trough-shaped clipping at an approximately 20° angle (dimensions: 60 mm x 17 mm, 0.4 mm deep) located centrally on the os frontale (Figure 3A). The cut is very sharp with slightly concave walls exhibiting fine, parallel, and slightly curved scratch marks at right angles to the lesion. The medial part of the wall is partially broken away due to later damage. There is no flaking, feathering, chattering, or peeling, but there are scoop defects toward the medial and posterior parts. The lesion is crossed by lesions 3 and 5 at its posterior and anterior ends.

Evaluation: This is a peri-mortem slash wound likely inflicted by a right-handed attacker of similar size in a standing position, striking from the front with a horizontal, whipping swing and a flick of the wrist at high velocity. The used weapon probably had a very sharp, light, and somewhat flexible blade (saber). This injury was neither lethal nor disabling.

Lesion 2 (CL2):

Description: A superficial, diagonal sharp cut from the left lateral down at a 30° angle (dimensions: 35 mm x 12 mm), extending through the left os zygomaticum, cleanly shearing off its lateral part and superficially glancing the maxilla's left alveolar rim, cutting and breaking off the left upper molars near the roots (Supplementary Figure 7). There is slight chattering along the superior defect margin but no flaking, feathering, scoops, or peeling. Sagittal cracks through the maxilla and the palatine process - following structural weaknesses along the roots of teeth 12 and 22, as well as through the frontal process of the maxilla - are observed.

Evaluation: This is a perimortem chop/slash wound inflicted from the front by a right-handed attacker of similar size in a standing position, striking diagonally inward with medium-high velocity. The weapon hada very sharp and thin edge. Though not immediately lethal, this severe facial injury was painful, disorienting, and impairing.

Lesion 3 (CL3):

Description: A steeply penetrating sharp blow at a flat diagonal angle (approximately 30°) across the lower frontal bone (lengths: 75 mm; depth: 19 mm), ending in lesion 1 (CL1) and breaking the right zygomatico-frontal junction (Figure 3A). Vertical cracks from the inferior cut margin may be results from unlodging the blade. The inferior margin is roughly cut and partially broken with flaking; there is no feathering, scoops, or peeling, but chattering along the defect margin.

Evaluation: This is a perimortem chop wound inflicted by a right-handed attacker of similar size from the right side of the victim in a standing position. The weapon was a moderately keen, long, straight, heavier blade (longsword). Though not immediately lethal, this wound was certainly life-threatening and staggeringly impactful.

Lesion 4 (CL4):

Description: A penetrating sharp blow at an acute angle to the left os parietale, close to and parallel to the coronal suture (cut lengths: 42 mm and cut length with breakage: 67 mm; width 9 mm; depth: 15 mm; angle: 20°) (Figure 3B). The lesion has a sharply cut posterior margin, while the anterior wall shows breakage, resulting in the loss of a narrow wedge of bone. The posterior margin shows flaking but no feathering, scoops, or peeling. Stress fractures towards the left orbita, sagittal suture, and along the left side of the coronal suture were caused by unlodging the blade.

Evaluation: This is a perimortem chop wound inflicted by a right-handed attacker of similar size or taller from the left side of the victim in a standing position. The weapon was a moderately keen, long, straight, heavier blade (longsword). Though not immediately lethal, this wound was certainly life-threatening and likely to stun and knock down the victim.

Lesion 5 (CL5):

Description: An extensive (dimensions: 50 mm x 33 mm) low-energy blunt force trauma at the hat brim line (HBL) level of the posterior half of the right os parietale/os temporale. This resulted in three radial fracture lines widening and diverging through the diploe and ending at sutural lines, with avulsions of the tabula interna due to deformation stress. The interior damage was only visible in CT images due to being covered by prior repairs. Vertical cracks through the lower os frontale, as described in lesion 3, could also be caused by bending stress from this impact.

Evaluation: This is a perimortem blunt trauma. The absence of a shaped impression or partial penetration suggests the injury was likely caused by a fall to the right side, followed by a hard impact of the head on the ground, although a kick to the side of the head with the victim kneeling or lying on the ground is also possible. The impact was probably forceful enough to cause dizziness or unconsciousness.

Lesion 6 (CL6):

Description: A deeply penetrating long diagonal sharp cut (length: cca. 100 mm; depth: 15 mms; skull vault thickness: 7 mm) through the upper part of the os frontale, cleanly severing a (lost) triangular bone segment between lesions 1, 3, and 6 (Figure 3A). The cut ends at lesions 3 and 4. The remaining superior wall is perpendicular to the lesion and smooth, with no flaking, feathering, scooping, or peeling, but slight chattering along the margin of the tabula interna. Slight damage in the form of torn layers in the cut wall due to compression is observable.

Evaluation: This is a perimortem chop wound inflicted from above with high velocity and force by a right-handed attacker in a higher position, likely standing over the downed victim slightly to the left. The weapon was a very sharp blade (saber). Though not immediately lethal, the injury was certainly life-threatening.

Lesion 7 (CL7):

Description: A sharply chop lesion (length: cca. 45 mm; width: 12 mm and with breakage on the maxilla: 22 mm), penetrating the right orbita at an almost vertical angle from the right lateral lower part, breaking off a bone chip from the maxillary orbital margin (Supplementary Figure 8A-B).

Evaluation: This is a perimortem chop wound inflicted with a wide, sharp blade (longsword). The victim's head was likely rolled to the left side. The attacker was standing above the victim towards the right side. This wound appears to be a deliberate mutilation or affective mangling of the defenseless victim’s face. The strike would have destroyed the eyeball.

Lesion 8 (CL8):

Description: A sharply cut small vertical lesion, cleanly cutting off the right lateral rim of the nasal aperture at an oblique angle, coming from the front/right side (length: 21 mm) (Supplementary Figure 8B). The cut margin shows no flaking, feathering, scoops, or peeling, but slight microcurvature of the exterior rim.

Evaluation: This is a perimortem stab/slash wound inflicted with a narrow, very sharp blade, possibly a saber, dagger, or knife. The victim's head was likely rolled to the left side, or the attacker used the left hand. This wound again appears to be a deliberate mutilation or affective mangling of the defenseless victim’s face.

Lesion 9 (CL9):

Description: A sharply cut small horizontal lesion (length: 20 mm), penetrating the left orbita at a steep angle from the medial upper corner part, coming from the front/slightly right side.

Evaluation: This is a perimortem stab wound inflicted with a narrow, sharp blade, possibly a saber, dagger, or knife. The victim's head was likely rolled to the left side, or the attacker used the left hand. This wound appears to be a deliberate mutilation or affective mangling of the defenseless victim’s face. The stab probably destroyed the eyeball, with lethality depending on the depth of penetration.

Additional Remarks

Surprisingly, the mandible displays no injuries corresponding to the multiple wounds inflicted on the cranium. The diagonal and irregular crack and gap in the mentum lack connecting pieces, preventing observation of original wound faces. A small post-mortem scratch is visible below the left foramen mentale.

The blow causing cranial lesion 2 smashed through the upper molars on the left side but did not damage the lower molars or mandible, which is geometrically difficult to explain. However, the complex combination of distance, positioning of the combatants, their movements, and residual force of the blow could account for this unexpected result.

Alternatively, the mandible might not originally belong to the victim and could have been added later to complete the find, either during interment, excavation, or reconstruction, intentionally or by mistake. The mandible fits moderately well, but the imperfect occlusal fit of the remaining molars and the proportionally longer lower ramus support this suspicion. So far, no explicit evidence such as separate DNA profiling is available.

*Supplementary Information 1.2. Postcranial Lesions*

(in order of probable infliction sequence)

Lesion 1 (PL1):

Description: A fine, superficial cut measuring 8 mm in length with slightly raised margins (micropeeling) is observed on the anterior surface of the right clavicle, 66 mm from the sternal end (Supplementary Figure 9).

Evaluation: This lesion is identified as a perimortem slash wound. The cut is nearly vertical and downward, indicative of a glancing, less forceful strike made from the front with a sharply edged blade. Flaking, feathering, chattering, or scoop defects are absent, slight marginal micropeeling is recognizable. The blade's tip might have damaged the subclavian artery and brachial plexus if the blade tilted and slid forward, but the steep angle of the cut makes this improbable. The weapon used was likely a saber.

Lesion 2 (PL2):

Description: A superficial, 19 mm long cut along the axis of the left clavicle, 28 mm from the sternal end (Supplementary Figure 10). The incision is tangential, impacting the clavicle at a diagonal angle from the lower anterior region, breaking off a small bone chip along the upper edge of the cut.

Evaluation: This is a perimortem superficial slash wound from a strike that came nearly horizontal in an inward arc, originating from the frontal right side. The fine cut mark suggests the very keen blade of a saber; the wound was probably only superficial.

Lesion 3 (PL3):

Description: A straight, tangential cut approximately 120 mm in length is observed along the spine of the right scapula, with the margin cleanly clipped and a strip of bone missing (Supplementary Figure 11). The lesion shows no flaking, feathering, peeling, or scoop defects, but exhibits slight marginal chittering.

Evaluation: This perimortem, vertical slash wound is consistent with a sharp blade wielded from the right side by a right-handed individual. The victim may have been standing or lying on the ground. The injury is extensive but rather superficial, likely inflicted by a longsword.

Lesion 4 (PL4):

Description: A fine, superficial and 14 mm long horizontal cut is present on the lateral-posterior aspect of the tuberositas deltoidea of the right humerus (Supplementary Figure 12). The cut features slight marginal chittering and flaking along both edges.

Evaluation: This is a perimortem chop wound; the kerf indicates an only slightly angled strike direction from distal to proximal from the right side, probably by a right-handed attacker in a standing position against the slightly raised arm of the victim. The microdamage to the cut walls indicates a moderately keen and heavier blade (longsword). The damage to the lateral head of the triceps brachii probably resulted in disabilities in movement of the right arm.

Lesion 5 (PL5):

Description: A very deep, sharply angled cut on the right ulna, measuring 18 mm in length, 13 mm in width, and 5 mm in depth, is located nearly at mid-shaft on the inferior surface (Supplementary Figure 13). The proximal wall is cleanly cut with slight marginal chittering and micropeeling, while the distal end is irregularly fractured with a missing wedge-shaped bone fragment. The kerf is straight, broad, and deep.

Evaluation: This perimortem backhand chop wound, inflicted from the front by a right-handed attacker in a standing position against a raised and bent right arm (indicative of a parry attempt), is likely the result of a saber. The nick on the interior margin of the right radius (Lesion 5b) may be associated with this injury (Supplementary Figure 14). The injuries caused by the strike sequence 5, 6, and 7 probably left the right forearm and hand disabled and bleeding.

Lesion 5b (PL5b):

Description: A superficial nick on the mid-shaft medial border of the right radius (Supplementary Figure 14).

Evaluation: This lesion is likely an extension of Lesion 5 (PL5).

Lesion 6 (PL6):

Description: A deep cut on the right ulna, similar in angle to Lesion 5, located 14 mm more distal, with a depth of 1.5 mm (Supplementary Figure 13). The proximal wall is cleanly cut with slight marginal chittering, while the distal end is irregularly fractured with two semicircular bone flakes missing. The kerf is straight, narrow, and deep.

Evaluation: This perimortem injury resembles Lesion 5 (PL5) but is less deep and at a steeper angle, suggesting a quick follow-up strike with the saber. The associated nick on the right radius (Lesion 6b - PL6b) is likely related.

Lesion (PL6b):

Description: A superficial nick on the mid-shaft medial border of the right radius, slightly more distal than Lesion 5b (PL5b) (Supplementary Figure 14).

Evaluation: This lesion is likely an extension of Lesion 6 (PL6).

Lesion 7 (PL7):

Description: A deep cut into the right ulna, oriented crosswise to Lesions 5 and 6, located 14 mm distal to Lesion 6, with a depth of 0.9 mm (Supplementary Figure 13). The proximal wall is cleanly cut with feathering, likely due to a rebounding blade. The cut shows a steep angle and a large bone flake tangentially removed. The kerf is straight, narrow and deep and indicates a strike angle much lower than in lesions 5 and 6.

Evaluation: The characteristics of this lesion, combined with those of Lesions 5 and 6, suggest a rapid sequence of strikes against the victim´s raised right arm, with only minor adjustments in positioning and strike direction.

Lesion 8 (PL8):

Description: A very deep diagonal downward cut on the lateral side of the left tibia, 25 mm below the tuberositas tibiae (Figure 4C). Measuring 35 mm in length and 12.6 mm in depth, the superior cut wall is sharply defined with slight chittering, but without flaking or feathering. The inferior wall shows rough breakage due to blade dislodgement.

Evaluation: This perimortem chop wound is from the front or left side by a right-handed attacker standing over the victim who was lying on the ground on the right side, with the leg pulled up to shield the body. The strike also affected the left fibula (Lesion 8b - PL8b). A distinction of blade type is not feasible based on the cut mark characteristics. The injury probably severed the common fibular nerve just above its branching.

Lesion 8b (PL8b):

Description: A steep-angled diagonal cut severing the proximal part of the left fibula, with irregularly sharp superior cut walls exhibiting slight chittering but no flaking or feathering (Supplementary Figure 15). The blow penetrated the bone and became lodged in the tibia.

Evaluation: This lesion is associated with the strike that impacted the left tibia (Lesion 8 - PL8) (Figure 4C), likely inflicted from the left side by a right-handed attacker.

Lesion 9 (PL9):

Description: A deep, steeply angled diagonal cut on the lateral side of the left tibia midshaft, measuring 16 mm in length and 5.2 mm in depth (Figure 4C). The superior cut wall is sharply defined without chittering, flaking, or feathering. The kerf is deep and somewhat irregular, the inferior cut wall broke off roughly by dislodging the blade, with a large flake removed.

Evaluation: This perimortem chop wound is consistent with a strike from the left side by a right-handed attacker standing over the victim who was lying on the ground on the right side, with the leg pulled up to shield the body. The lesion characteristics suggest a heavier and moderately keen blade (longsword).

Lesion 10 (PL10):

Description: A slightly oblique, long, and deep cut (24 mm in length and 3.8 mm in depth) on the left femur's antero-lateral side just below the trochanter major with a trailing scratch towards the lateral (Figure 4D). The proximal cut wall is slightly concave without flaking, chittering, or feathering, while the distal wall shows significant breakage with a large lunate fragment missing. The kerf extends into a fine fracture line.

Evaluation: This perimortem forceful vertical chop from the front by a right-handed attacker again likely occurred against the victim lying on the ground and trying to shield himself with an upraised right leg. A longsword is the probable weapon.

Lesion 11 (PL11):

Description: A horizontal, long cut (21 mm in length and 1.8 mm in depth) on the left femur, antero-lateral side approximately mid-thigh (Figure 4D). The proximal wall is straight with minimal chittering but no flaking or feathering, while the distal wall shows shallow breakage from blade rebound, with a large rectangular flake missing. The kerf extends into a fine fracture line.

Evaluation: This perimortem fast vertical chop from the left side by a right-handed attacker likely occurred with the victim lying on the ground and attempting to shield himself with an upraised right leg. A longsword is the probable weapon. The strike may have severed the lateral circumflex femoral artery, leading to substantial hemorrhage.

Lesion 12 (PL12):

Description: An oblique, long cut (20.8 mm in length and 3.8 mm in depth) on the left femur, antero-lateral side mid-thigh (Figure 4D). The proximal cut wall is straight with slight chittering and flaking near the lateral end, while the distal wall shows steep-angled breakage with a long wedge-shaped fragment missing.

Evaluation: This perimortem forceful angled chop from the left side by a right-handed attacker likely targeted a victim lying on the ground and attempting to shield himself with an upraised right leg. The weapon used was probably a longsword.

Lesion 13 (PL13):

Description: A vertical tangential cut through the processus spinosus of the lumbar vertebra L2, with the posterior half of the processus spinosus sheared off.

Evaluation: This perimortem slash wound was inflicted by a right-handed attacker standing over the victim, who was lying on their right side. The victim was likely in a curled position, exposing the lumbar processus spinosus. The sharp edge of the weapon suggests a saber.

Lesion 14 (PL14):

Description: A horizontal cut through the processus spinosus of thoracic vertebra T7, removing the lower part (Figure 4A). The cut did not penetrate the spinal canal but ended on the articular surfaces of thoracic vertebra T8, with slightly greater depth on the left side.

Evaluation: This perimortem chop wound was inflicted from the front/left side by a right-handed attacker standing over the victim lying on their right side. The weapon was likely a saber.

Lesion 15 (PL15):

Description: A nearly horizontal stab from behind and slightly to the left, penetrating the spinal canal between thoracic vertebrae T10 and T11 (Figure 4B). The point entered the vertebral body of T11 from behind, with very sharp and clear margins.

Evaluation: This perimortem stab wound is consistent with a symmetrical, flat lenticular, two-edged weapon. The stab breadth is 28.2 mm, the weapon was likely a longsword. The severing of the spine would have resulted in paralysis of the lower body.

Lesion 16 (PL16):

Description: A diagonal deep cut through the proximal end of the left ulna (Supplementary Figure 16) from proximal to distal direction, nearly severing it, with incidental grazing of the proximal end of the radius (Lesion 16b - PL16b). The cut walls show post-mortem damage.

Evaluation: This perimortem slash wound was inflicted from a vertical, slightly inward swing from the left front side by a right-handed attacker. The victim's left arm was likely raised in a blocking position. The weapon was possibly a saber.

Lesion 16b (PL16b):

Description: A diagonal deep grazing cut into the proximal end of the left radius, from distal to proximal direction. An indentation from lodging and unlodging the blade is visible (Supplementary Figure 17).

Evaluation: This perimortem slash wound, inflicted from a vertical, slightly inward swing from the front by a right-handed attacker, indicates a blocking position of the left arm. The weapon used was possibly a saber. Position and direction of force make a connection to Lesion 16 (PL16) (Supplementary Figure 16) probable.

Lesion 17 (PL17):

Description: A steep-angled short, but forceful cut into the tuberositas radii of the left radius (Supplementary Figure 17) probably from proximal to distal.

Evaluation: Position and direction of force make PL16 as common cause unlikely; rather, it represents a separate injury. The situational sequence with regard to PL16 and PL16b is unclear; it could represent an earlier more superficial hit from the same attacker against the not fully raised and more outstretched arm.

**Supplementary Information 2. Weapon characteristics**

All observed lesions are caused by chops, slashes or stabs inflicted by metal blade implements that penetrated soft tissue and damaged underlying bone. Following the criteria of kerf length and width, cut wall angle and profile, surface damage like feathering, chittering or flaking, and kerf bottom profile, the use of two different kinds of long blades could be derived:

One was probably a comparatively light and quick, very sharp blade used for slashing attacks, while the other type was heavier and slower, with lower sharpness and used for chopping (chopping means a strike perpendicular to the surface of the target, cleaving vertically through tissue, while slashes are also drawn along the blade`s long axis, making a cutting move).

With respect to weaponry used in the region in the late 13th century AD, the first weapon could be identified as a Cuman-style saber, while the second type was a double-edged longsword of Oakeshotte´s Type XII or XIII.

Saber-type blades (curved backswords) were introduced into the region of today's Hungary by the Avars as a slashing weapon well suited to mounted combat, and in use in slightly varying types later by Magyar, Cuman, Mongol and Ottoman invaders. They were ca. 80-90 cm long, 3-4 cm wide and 3-4 mm thick at the back, slightly curved with a sharpened outer edge; the tip's back was also sharpened for about a quarter of the blade`s length, the blade´s profile was cuneiform (flat grind to the edge). Total weight was around 600 to 700 g. The effectiveness of the saber was based on swift slashes and easy changes of attack direction, although long stabs were possible.

Longswords, derived from the Merovingian and Carolingian spatha-type symmetric double-edged swords, were predominantly used in the German regions. With their longer and broader blades (ca. 85-100 cm long, 4-6 cm wide) they were heavier than sabers, but thanks to the pommel-counterweight balanced and not as cumbersome as often envisioned. Still, their total weight including pommel and quillons was usually 1300-1500 g. They were mainly used for heavy chops and short stabs. To better resist contact with metal armor, which was more common in Western European warfare at that time, the edge was not kept at maximum keenness.

The blade's profile, to better resist the impact of heavy chops and stabs against hard armor, was symmetrical, lenticular (convex grind to the edges) and fullered.

Both weapons differ in the way they are wielded, and in the damage characteristics they inflict on bone; following Lewis' categories, the longsword belongs to the broadsword class and the saber to the scimitar class.

#### Supplementary Information 3. Forensic evaluation and situational reconstruction

##### *Supplementary Information 3.1. Forensic traumatological assessment*

Based on the completeness and preservation of the skeleton, as well as the clarity of all observable perimortem lesions (9 on the skull and 17 on the postcranial bones), the relevant forensic questions could be addressed with a high degree of confidence. All injuries exhibited by the examined individual were sustained perimortem within a very short time frame and were undoubtedly inflicted during a single incident of interpersonal violence, with some even overlapping. No relevant prior injuries were present on the skeleton, and all lesions were perimortem and inflicted with intent. The injuries observed were either caused by direct attacks or were defensive wounds incurred while attempting to fend off the attacks. Even the fractures on the right side of the skull — likely resulting from a hard fall — were directly caused by the violent encounter rather than an accident. The remaining lesions were clearly inflicted by implements specifically designed for interpersonal conflict, indicating an intent to harm and likely to kill.

The varying characteristics of the damage marks and the diverse vectors of attack suggest that at least two, possibly three assailants were involved: one attacking from the front, another from the left side, and a third from the right side. The intent was likely to prevent the victim from escaping and to ensure a decisive and quick outcome through overwhelming force. Therefore, the situation was not a duel but an assassination.

While certain characteristics of skeletal lesions can help differentiate the types of blades used, these criteria are not entirely unambiguous. Cut margins are influenced by weapon characteristics such as blade curvature, thickness, profile shape, and weight—traits defining different weapon categories. However, individual factors such as blade sharpness, attacker strength and speed, specific strike angle, or body protection worn by the victim can significantly influence the appearance of such lesions. Additionally, the damage patterns on bones with a hard and thick cortex differ from those on more spongy bone parts that do not resist surface penetration as well.

Despite these variables, the observed lesions suggest that at least two different types of blades were used. One type, of lighter weight, was employed repeatedly in quick slashing blows, had a very keen and thin edge with a wedge-shaped cross-section, and can probably be defined as a saber-like backsword. The other type, heavier and slower, was used primarily for chopping and stabbing, with a less sharp edge and a symmetrical flat lenticular blade. Considering the weapons used in the late 13th century in the region, these weapons could probably be identified as a Cuman-style saber and a western longsword of Oakeshott Types XII or XIII.

The majority of the injuries were inflicted by an attacker positioned in front of the victim, with a few strikes coming from the left and right sides, and none from behind. This pattern suggests an open confrontation, aligning with historical accounts. The primary attacker, likely Henrik Kőszegi, can thus be identified as the assailant from the front wielding the saber.

Overall, the pattern and severity of the attacks, as evidenced by the injuries, indicate an intent to kill. The extent of the violence and the mutilating injuries to the victim's face suggest a high degree of emotional involvement, such as rage and hatred. Nonetheless, premeditated murder cannot be ruled out; the altercation could have been planned and provoked to provide justification for the attack, and the coordinated assault by multiple attackers implies some level of pre-planning.

*Supplementary Information 3.2. Reconstruction of the situation*

The presence of lesions inflicted by at least two distinct bladed weapons, likely a saber and a longsword, suggests the involvement of at least two different assailants. The reconstructed directions and positioning of the wounds further support the plausibility of simultaneous attacks by three individuals. Based on the sequence and characteristics of the attacks, it appears that the primary assailant, wielding the saber and positioned frontally, delivered the majority of the lethal strikes, while the secondary assailants, wielding longswords and flanking the victim on the left and right, assisted in the attack.

To accurately reconstruct the phases of the assault, it is necessary to sequence each attack in the correct order. This is most reliably achieved through visible overlaps, such as one cut crossing an earlier one or a fracture line stopping at an earlier fissure. In this case, most lesions are independent, necessitating the application of less reliable criteria. One such criterion is proximity: in the dynamic situation of a melee involving multiple attackers, it is improbable that the same body part would be struck independently in the same manner again. More often, adjacent similar wounds are inflicted in quick succession by follow-up strikes. Another assumption in a lethal conflict is the severity of wounds: initially, the defender is more successful in avoiding hits (if not caught unaware), but as fatigue and injuries accumulate, subsequent attacks become more effective. These aspects were considered in reconstructing the murder scene.

The clearly defined sharp cut marks on the bones, some very deep, and the absence of crushing or compression damage indicate that the victim was not protected by sturdy body armor. The head injuries, in particular, would not have been possible if the victim had been wearing a helmet. Furthermore, the parry injuries on both forearms suggest desperate unarmed defense attempts, indicating that the victim may have been surprised by the initial attacks, caught without a weapon, or quickly disarmed. The high number of hits suggests that the attackers were able to execute a flurry of blows against a mostly defenseless target.

A plausible reconstruction of the scene involves Kőszegi confronting Duke Béla in an argument, suddenly drawing his saber and delivering a series of quick slashes to the head and upper body (CL 1-2, PL 1-2). Meanwhile, assailants 2 and 3 flanked the victim and attacked him with their longswords (CL 3-4, PL 3-4), while the victim attempted to fend off further strikes from the front, resulting in deep saber slashes to the right forearm (PL 5-7). One of the head blows (likely head lesion 4 from the left side) struck the prince down to his right side, where his injured right arm could not effectively break the fall, causing him to crack his head hard on the ground. Dazed by the impact, the victim, lying on his back/right side, lifted his left arm and leg to shield himself against further strikes. Kőszegi then stepped over him, wildly slashing at the head (CL 6), arm (PL 16), and body (PL 13, 14), while assailant 2, positioned on the victim's left side, chopped at the leg and stabbed him in the back (PL 8-12, 15). The facial wounds (PL 7-9) were likely inflicted when the victim was no longer moving, with his head lolled to the left side and his face exposed to the attacks from the front and right side.

**Supplementary Information 4. Dietary patterns of the investigated young male based on the results of the dental calculus analysis**

In the years following Béla’s birth, Hungary's population began to recover from the food shortages caused by the Mongolian invasion. Like other medieval Central European nations, Hungary primarily cultivated C3 cereal crops. Alongside wheat and barley, rye (*Secale cereale*) and oats (*Avena sativa*) were also grown during this time. The C4 common millet (*Panicum miliaceum*) was another common grain. Typically, both grains and legumes were consumed as porridge. After intensive grinding, people could eat cereal grains in various forms: as baked or boiled foods, barley porridge, or baked barley pies.

Duke Béla's dental plaque matrix contained several gelatinized starches, which no longer exhibited birefringence under polarized light (Supplementary Figure 5). However, the layers and crystallization foci could still be recognized in some cases. Regarding the size, morphological and light refraction characters of the starch grains and the material of the gelatinized starch in the dental calculus were similar to the reference material of the boiled wheat grits and the baked wheat bread. In Duke Béla’s dental calculus samples, the shape of the starch granules did not clearly show features characteristic of any specific cereal type, yet wheat and barley appear to be likely candidates based on the size and shape of the starch grains (Supplementary Figure 6). Given the significant presence of long plant hairs, we can also infer a considerable amount of wheat consumption.

The signs of cooking include grooves that start radially from the crystallization focus of amyloplasts, and the triangular shape of the starch grain may suggest grinding (1). Intensely ground or pounded kernels, along with suspected yeast cells, indicate the consumption of fermented foods, suggesting that leavened wheat bread was likely eaten (2–4). Bread made from fermented dough was already consumed by the aristocracy in the 11th century, but it did not become widespread among common people until the 14th century, and even then, it was only eaten in small quantities (5). Domestic artifacts that clarify bread consumption were absent during this period. The presence of several pollen grains in the sample may suggest the consumption of honey or honey-infused foods. Though, recently, researchers have concluded that when assessing the origin of the residue, it is also important to consider that inhaled substances, like pollen, can be incorporated into tartar as well (6). Notably, numerous Ca-oxalate crystals were found in the calculus matrix (7). In addition to the three separate styloid- and druse-shaped crystals, a starchy tissue residue was also found, in which 5-6 crystal druses and their broken pieces were wedged. Calcium oxalate crystals are rarely preserved in archaeological calculus (7), and acidic sample preparation can dissolve them (8). Therefore, ancient calcium oxalate crystals found in calculus are rarely reported. Although many plant parts can contain Ca-oxalate, crystal druses and starch were more concentrated and occurred together, which is a characteristic of plant seeds rather than shoots. Identified based on our reference source material (labcode: ML207246) and Decke's definition key (9), the consumption of hazelnut seeds (*Corylus avellana* L.) cannot be ruled out.

### Supplementary Figures

**Supplementary Figure Legends**

Supplementary Figure 1. The skeletal remains of an adult male were found beneath the floor during the excavation of a 13th century Dominican monastery on Margaret Island, in 1914-1915. (source: photo by János Müllner; Budapest History Museum - Kiscelli Museum, Inv. nr. 10033.) (10)

Supplementary Figure 2. The skeletal remains of the investigated individual were found beneath the floor during the excavation

Supplementary Figure 3. The collagen fraction of bone samples for radiocarbon yielded conventional $^{14}C$ dates (n=7) between 874±19 BP and 922±17 BP, corresponding to a combined calibrated age range of cal AD 1050-1180 at 95.4% probability level. The youngest age was offered by the outermost layer of the bone, which is supposed to have been contaminated by some preservatives during the processing in the collections. The bioapatite date of 841±18 BP corresponds to a calendar age of AD 1170-1260.

Supplementary Figure 4. A-B: The left upper 1st premolar and 1st-2nd molars are fractured in the region of the tooth neck, presumably due to forced impact immediately before death. The plane of fracture of the two molars is the same as the plane of the cut of the zygomatic bone and the vestibular bony process of the upper jaw around those molars. C: Non dislocated fracture of the mesiobuccal root of the left upper first molar tooth

Supplementary Figure 5. Microremains recovered from the dental calculus (the opaline nature of a few images is due to the partly dissolving calculus matrix in HCl). (A) Gelatinous starch grains from Triticeae fruits and their polarized microscopic images. (B) and (C) Ground and cooked degraded starch grains and their polarized microscopic images. (D) Plant hairs. (E) Calcium-oxalate crystal. (F) The tissue of a cereal endospermium with bimodal-sized starch grains sticking into that. (G) and (H) Pollens. (I) and (J) Spores and microcharcoal in the dental calculus matrix (arrows). (K) Cell from the pericarpium of a cereal fruit. (L) Phytolith. (M) Sponge spicule. (N) Supposed yeast cells integrated in the dental calculus matrix (note the budding morphology and the vacuolated structure). (O) Undefined microremain. (P) Supposed nematode remains. Scale bars are 10 μm except for (A, D, F) where 100 μm.

Supplementary Figure 6. Microremains recovered from the dental calculus with their comparing modern reference material pairs. (A) Cereal fruit cell from the dental calculus. (B) Wheat grit starch grain (reference). (C) Supposed endosperm cell from the dental calculus and (D) from wheat bread with blue discoloration (reference). (E) Trichome from the dental calculus and (F) from wheat bread (reference). (G) A ground starch grain from the dental calculus and (H) from boiled wheat grit. (I) Supposed yeast cell aggregate from the dental calculus and (J) from wheat bread (reference). Scale bars are 10 μm.

Supplementary Figure 7. The lateral part of the left zygomatic bone was cut off by a sharp weapon (A). The same cut superficially glanced the maxillary left alveolar rim, cutting and breaking off the left upper molars near the roots (B). CL2=Cranial Lesion 2. Detailed description of the perimortem lesions is available in Supplementary Information 1.1.

Supplementary Figure 8. Perimortem cut marks on the right (A) and left zygomatic bones and maxillae (B) caused by sharp weapons. CL2=Cranial Lesion 2; CL7=Cranial Lesion 7; CL8=Cranial Lesion 8. Detailed description of the perimortem lesions is available in Supplementary Information 1.1.

Supplementary Figure 9. A superficial cut mark is observed on the anterior surface of the right clavicle. PL1= Postcranial Lesion 1. Detailed description of the perimortem lesion is available in Supplementary Information 1.2.

Supplementary Figure 10. A superficial cut is observed on the left clavicle. PL2= Postcranial Lesion 2. Detailed description of the perimortem lesion is available in Supplementary Information 1.2.

Supplementary Figure 11. A straight, tangential cut can be seen along the spine of the right scapula, with the margin cleanly clipped and a strip of bone missing. PL3= Postcranial Lesion 3. Detailed description of the perimortem lesion is available in Supplementary Information 1.2.

Supplementary Figure 12. A fine, superficial horizontal cut is present on the lateral-posterior surface of the tuberositas deltoidea of the right humerus. PL4= Postcranial Lesion 4. Detailed description of the perimortem lesion is available in Supplementary Information 1.2.

Supplementary Figure 13. Two perimortem cut marks can be observed on the right ulna. A deep, sharply angled cut is located nearly at mid-shaft on the inferior surface (PL5) and another clear cut (PL6) is present 14 mm more distal from PL5. PL5= Postcranial Lesion 5; PL6= Postcranial Lesion 6. Detailed description of the perimortem lesions are available in Supplementary Information 1.2.

Supplementary Figure 14. Two superficial nicks on the mid-shaft medial border of the right radius (PL5b and PL6b) are likely extensions of PL5 and PL6. PL5= Postcranial Lesion 5; PL6= Postcranial Lesion 6; PL5b= Postcranial Lesion 5b; PL6= Postcranial Lesion 6b. Detailed description of the perimortem lesions are available in Supplementary Information 1.2.

Supplementary Figure 15. A diagonal strike severing the proximal end of the left fibula. PL8b= Postcranial Lesion 8b. Detailed description of the perimortem lesion is available in Supplementary Information 1.2.

Supplementary Figure 16. A cut through the proximal end of the left ulna from the dorsal side. PL16= Postcranial Lesion 16. Detailed description of the perimortem lesion is available in Supplementary Information 1.2.

Supplementary Figure 17. Two sharp lesions cut into the proximal end of the left radius. PL16b= Postcranial Lesion 16b, PL17= Postcranial Lesion 17. Detailed description of the perimortem lesions is available in Supplementary Information 1.2.

Supplementary Figure 18. $^{87}Sr/^{86}Sr$ values on different tissues of the subject, compared with local baselines, collected by Price et al. (11), Depaermentier et al. (12), Cavazzuti et al. (13)

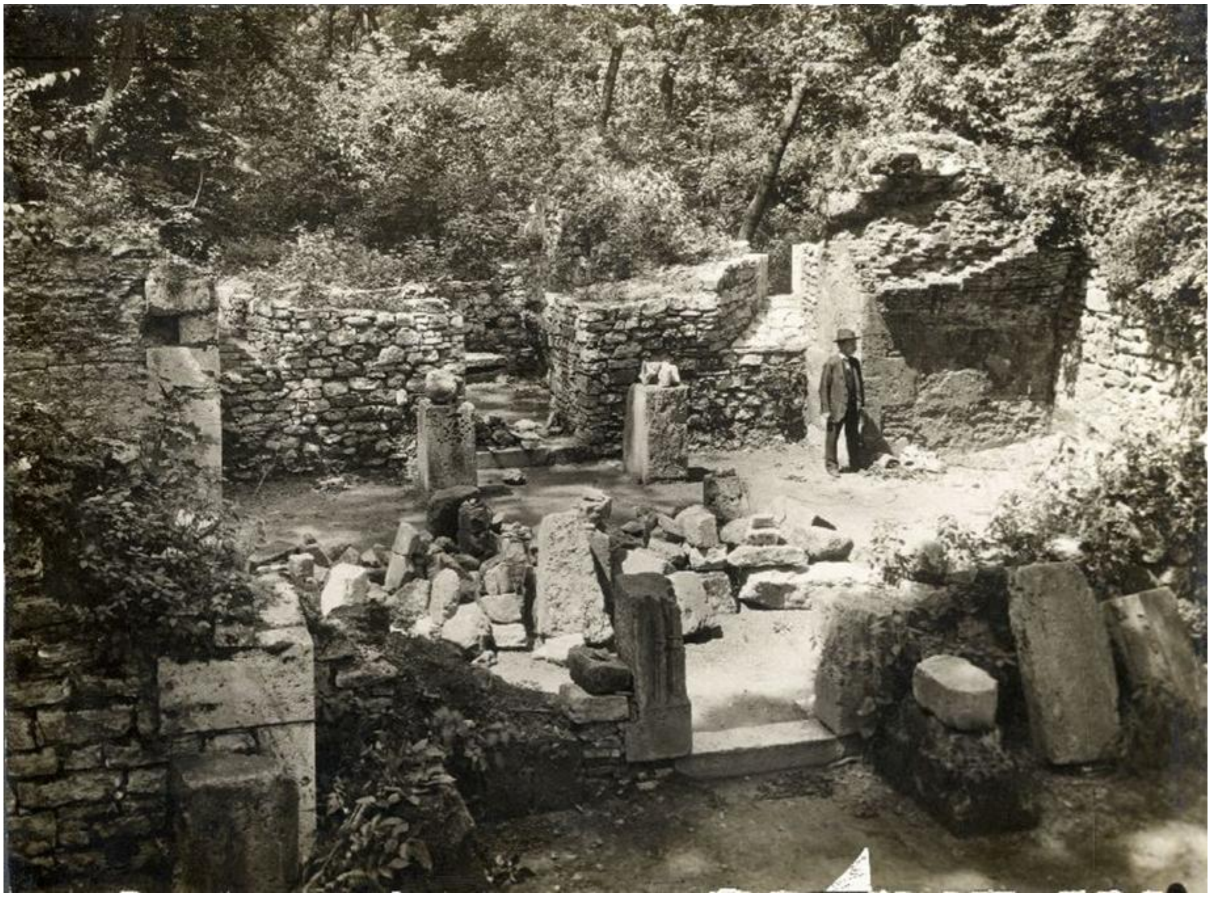

Supplementary Figure 1. The skeletal remains of an adult male were found beneath the floor during the excavation of a 13th century Dominican monastery on Margaret Island, in 1914-1915. (source: photo by János Müllner; Budapest History Museum - Kiscelli Museum, Inv. nr. 10033.) (10)

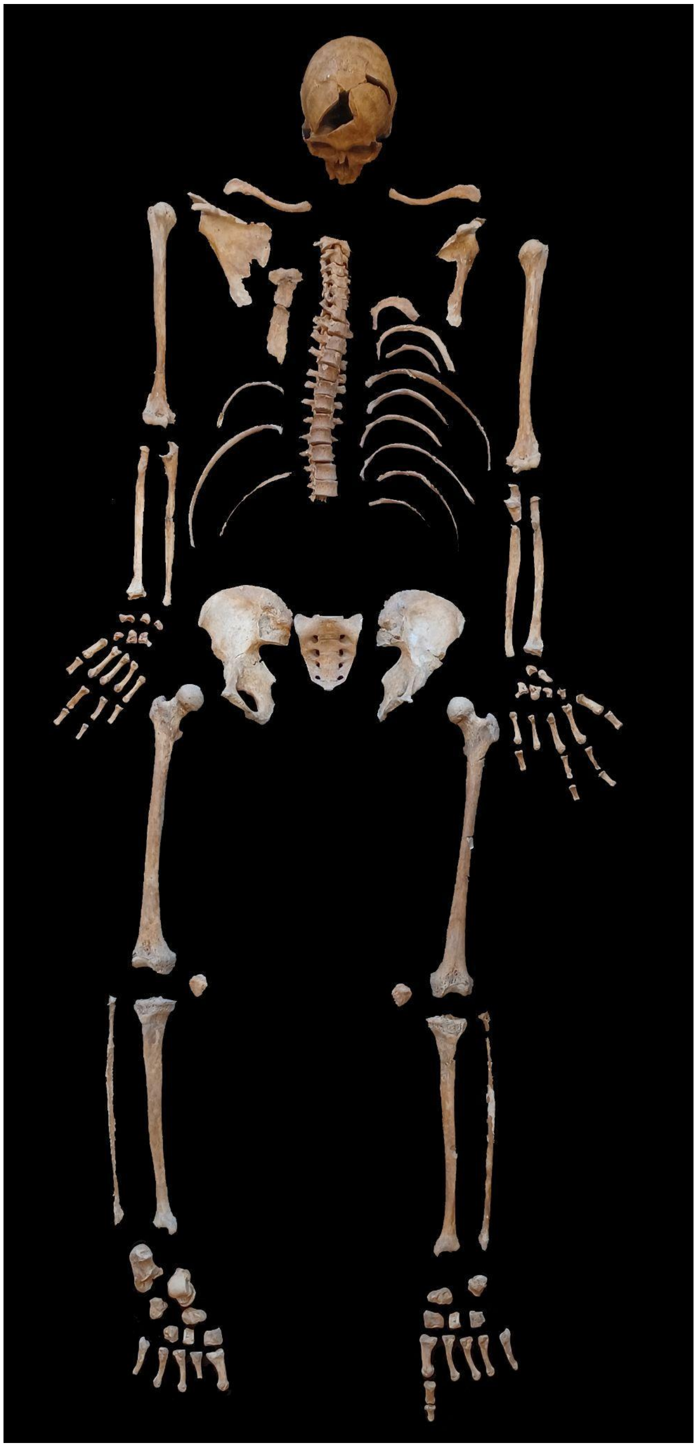

Supplementary Figure 2. The skeletal remains of the investigated individual were found beneath the floor during the excavation

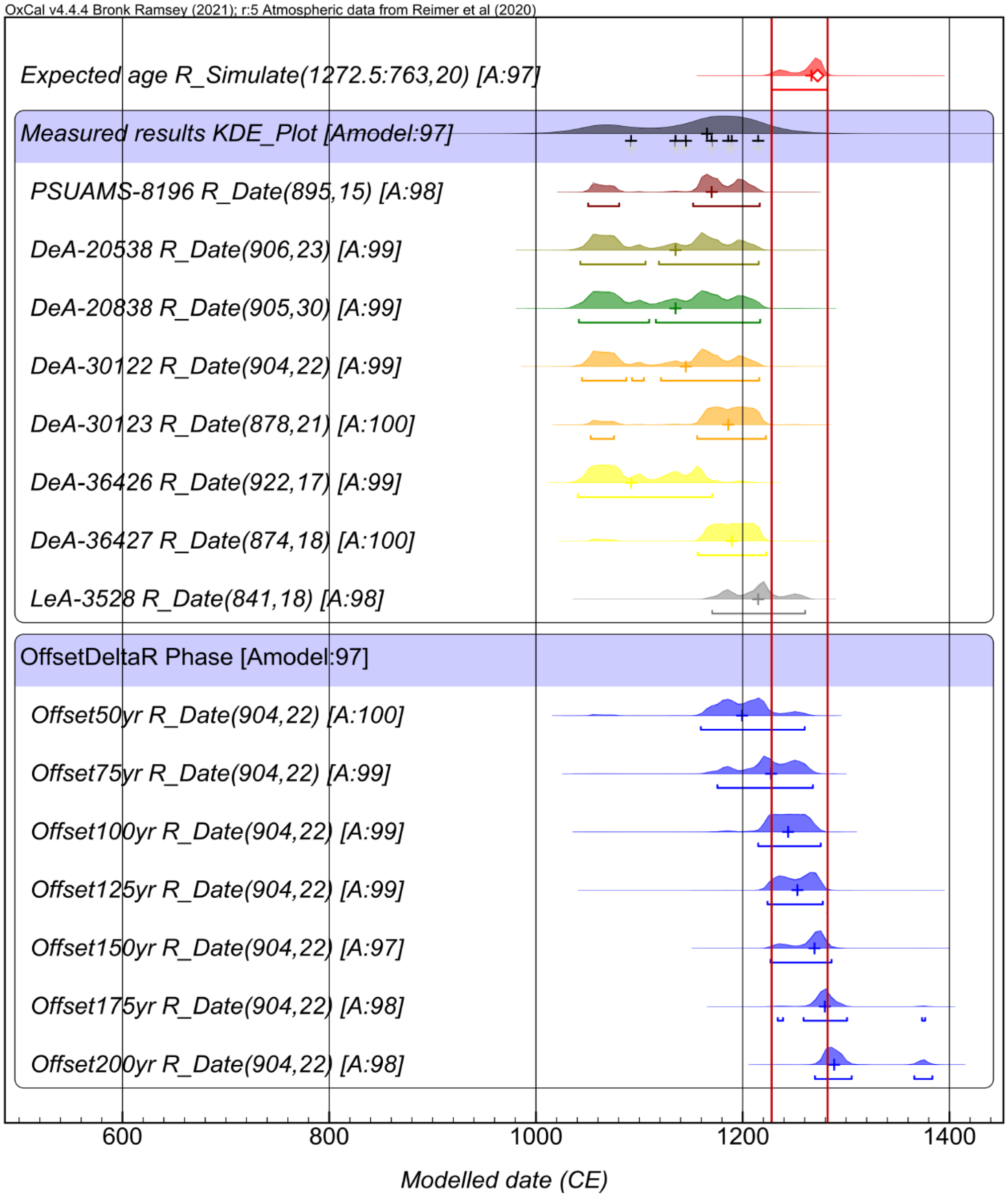


Supplementary Figure 3. The collagen fraction of bone samples for radiocarbon yielded conventional $^{14}C$ dates (n=7) between 874±19 BP and 922±17 BP, corresponding to a combined calibrated age range of cal AD 1050-1180 at 95.4% probability level. The youngest age was offered by the outermost layer of the bone, which is supposed to have been contaminated by some preservatives during the processing in the collections. The bioapatite date of 841±18 BP corresponds to a calendar age of AD 1170-1260.

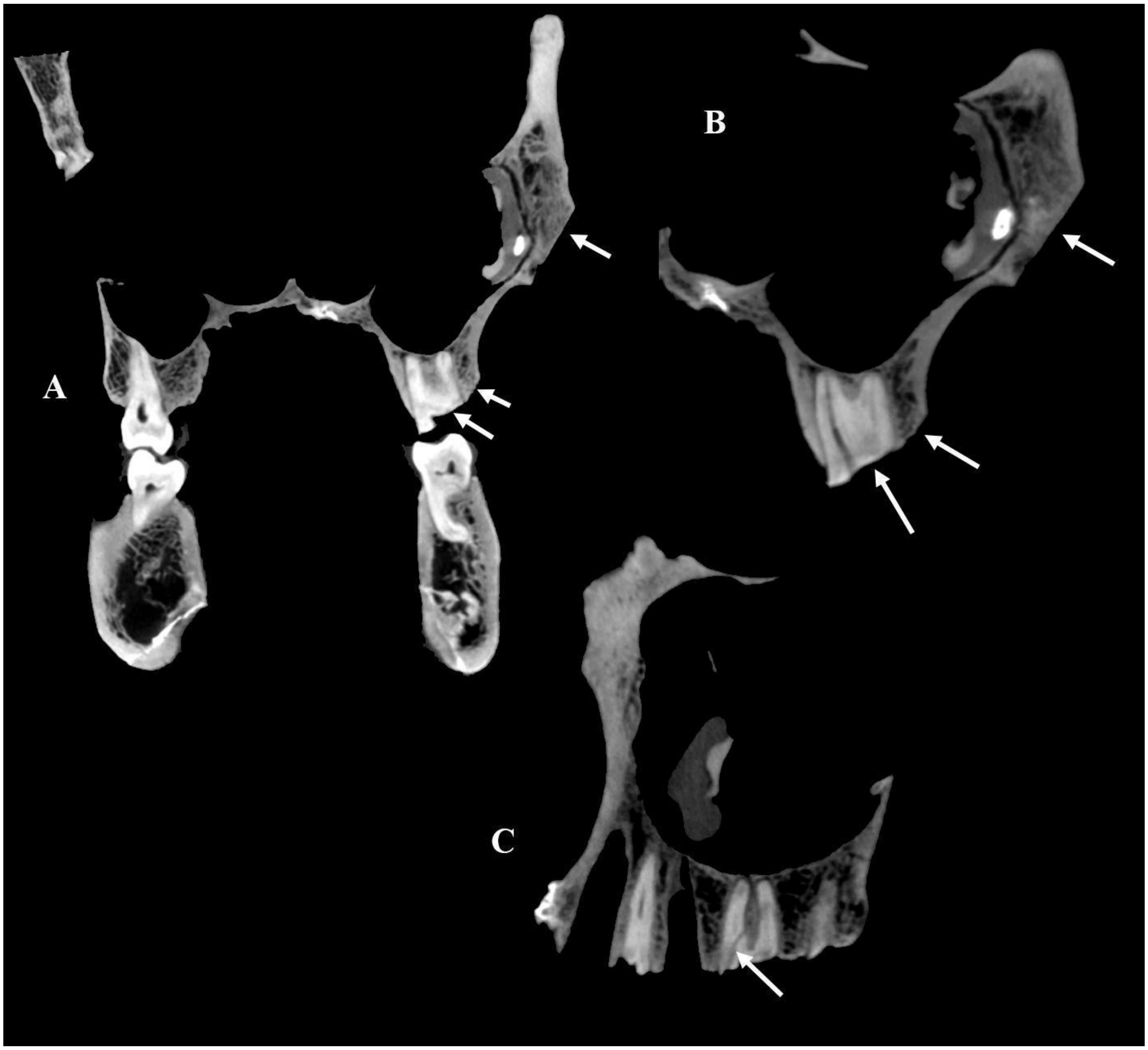


Supplementary Figure 4. A-B: The left upper 1st premolar and 1st-2nd molars are fractured in the region of the tooth neck, presumably due to forced impact immediately before death. The plane of fracture of the two molars is the same as the plane of the cut of the zygomatic bone and the vestibular bony process of the upper jaw around those molars. C: Non dislocated fracture of the mesiobuccal root of the left upper first molar tooth

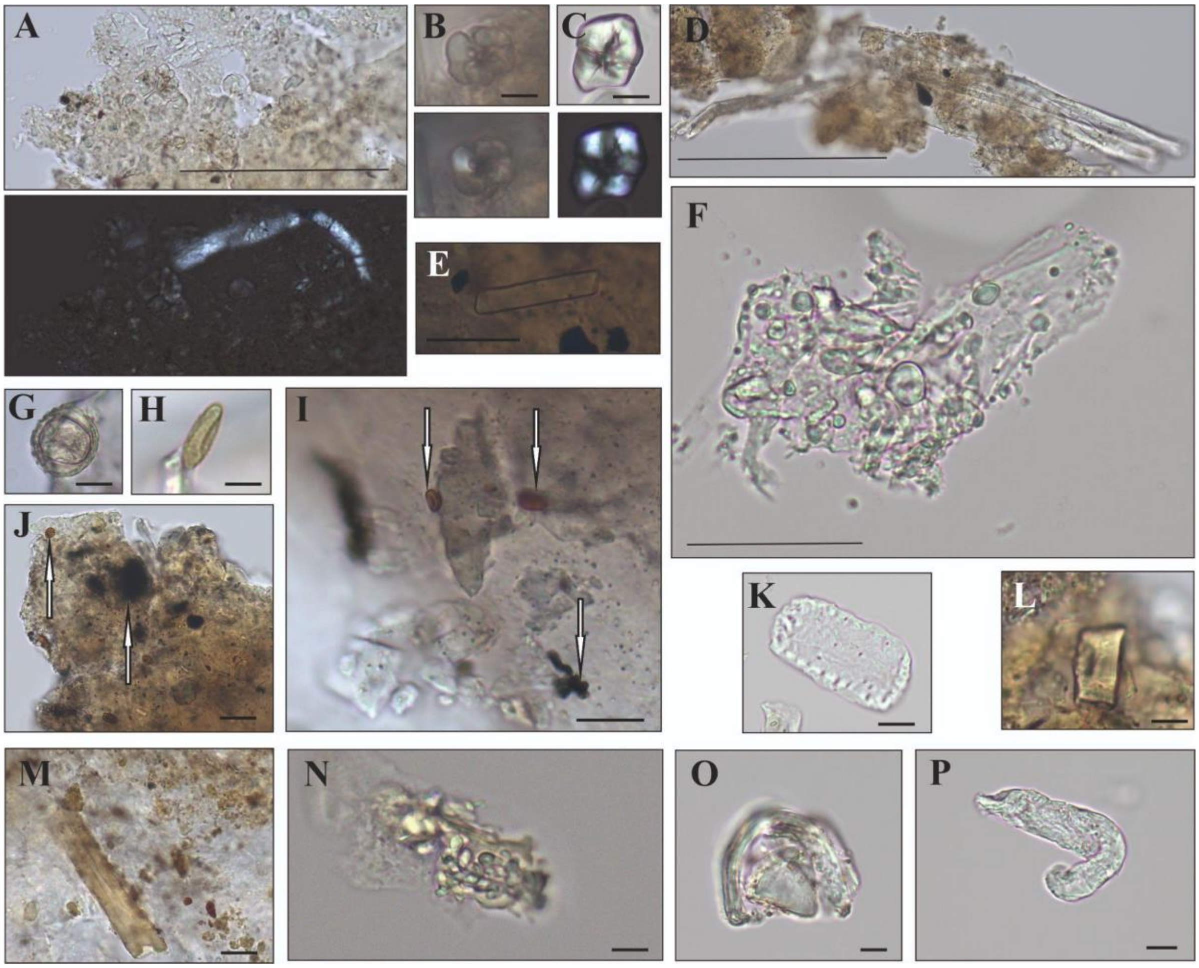


Supplementary Figure 5. Microremains recovered from the dental calculus (the opaline nature of a few images is due to the partly dissolving calculus matrix in HCl). (A) Gelatinous starch grains from Triticeae fruits and their polarized microscopic images. (B) and (C) Ground and cooked degraded starch grains and their polarized microscopic images. (D) Plant hairs. (E) Calcium-oxalate crystal. (F) The tissue of a cereal endospermium with bimodal-sized starch grains sticking into that. (G) and (H) Pollens. (I) and (J) Spores and microcharcoal in the dental calculus matrix (arrows). (K) Cell from the pericarpium of a cereal fruit. (L) Phytolith. (M) Sponge spicule. (N) Supposed yeast cells integrated in the dental calculus matrix (note the budding morphology and the vacuolated structure). (O) Undefined microremain. (P) Supposed nematode remains. Scale bars are 10 μm except for (A, D, F) where 100 μm.

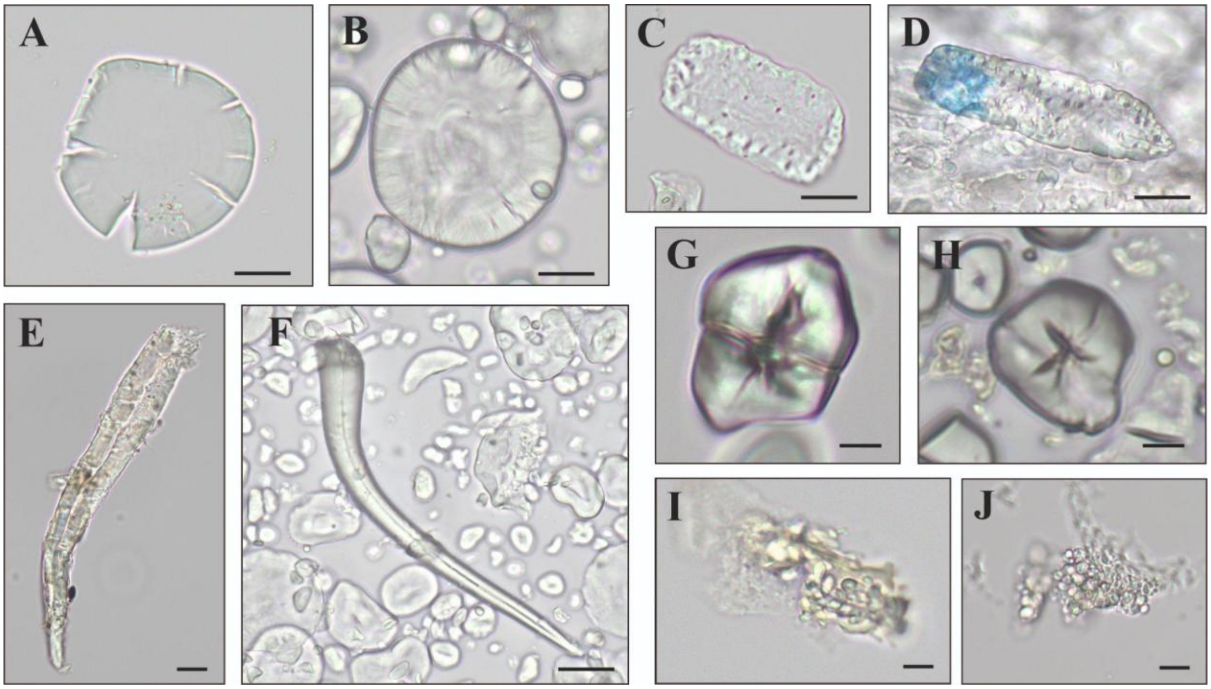


Supplementary Figure 6. Microremains recovered from the dental calculus with their comparing modern reference material pairs. (A) Cereal fruit cell from the dental calculus. (B) Wheat grit starch grain (reference). (C) Supposed endosperm cell from the dental calculus and (D) from wheat bread with blue discoloration (reference). (E) Trichome from the dental calculus and (F) from wheat bread (reference). (G) A ground starch grain from the dental calculus and (H) from boiled wheat grit. (I) Supposed yeast cell aggregate from the dental calculus and (J) from wheat bread (reference). Scale bars are 10 µm.

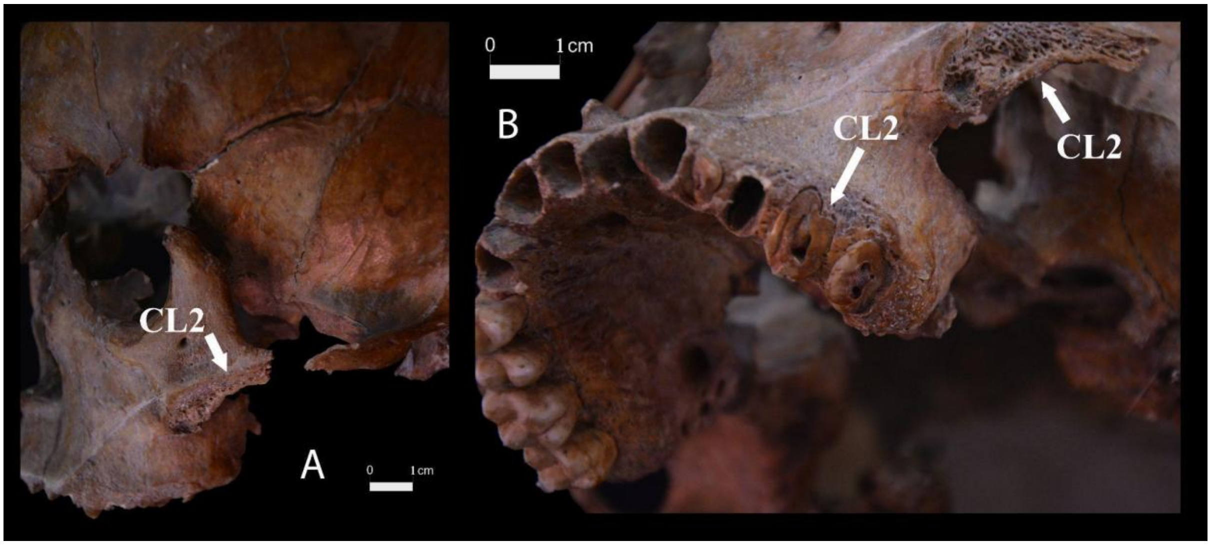


Supplementary Figure 7. The lateral part of the left zygomatic bone was cut off by a sharp weapon (A). The same cut superficially glanced the maxillary left alveolar rim, cutting and breaking off the left upper molars near the roots (B). CL2=Cranial Lesion 2. Detailed description of the perimortem lesions is available in Supplementary Information 1.1.

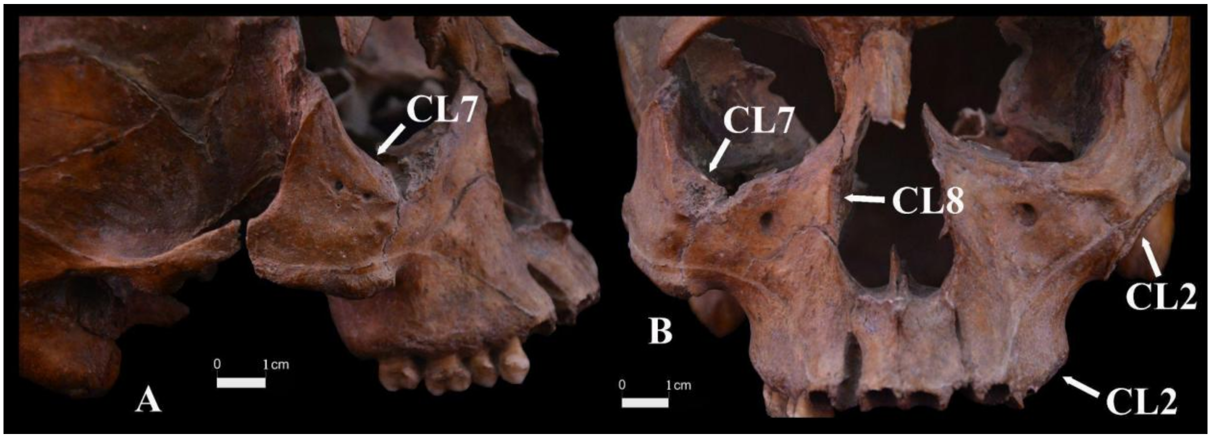


Supplementary Figure 8. Perimortem cut marks on the right (A) and left zygomatic bones and maxillae (B) caused by sharp weapons. CL2=Cranial Lesion 2; CL7=Cranial Lesion 7; CL8=Cranial Lesion 8. Detailed description of the perimortem lesions is available in Supplementary Information 1.1.

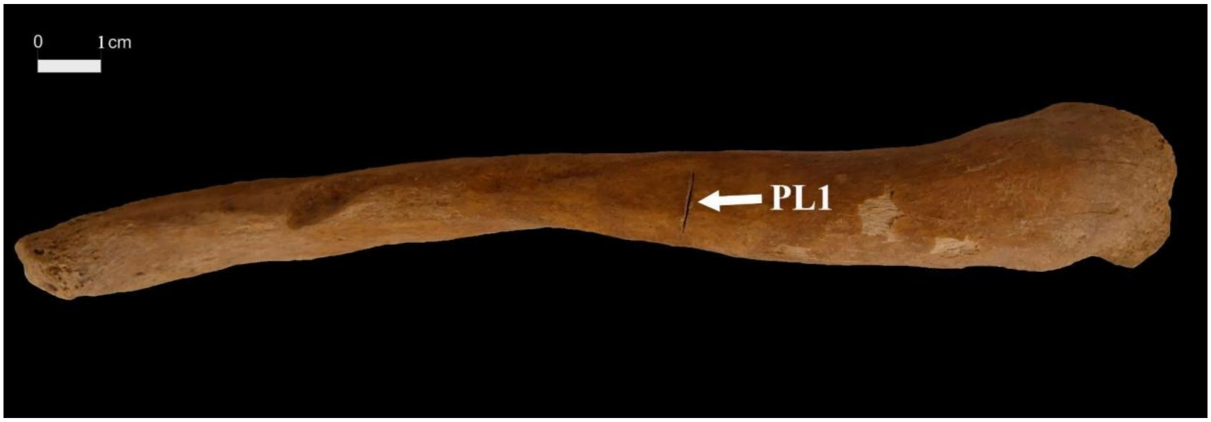


Supplementary Figure 9. A superficial cut mark is observed on the anterior surface of the right clavicle. PL1= Postcranial Lesion 1. Detailed description of the perimortem lesion is available in Supplementary Information 1.2.

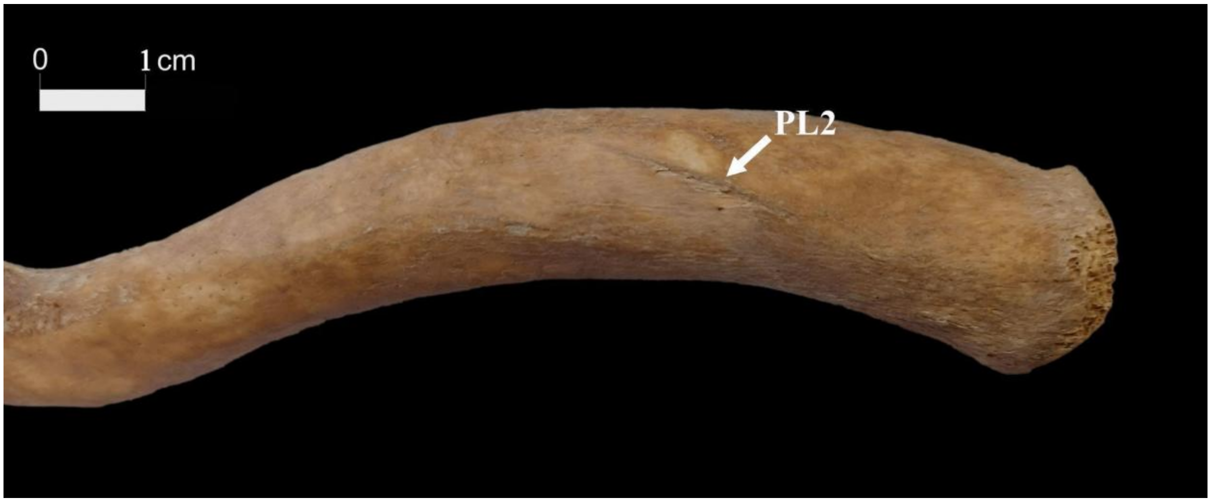


Supplementary Figure 10. A superficial cut is observed on the left clavicle. PL2= Postcranial Lesion 2. Detailed description of the perimortem lesion is available in Supplementary Information 1.2.

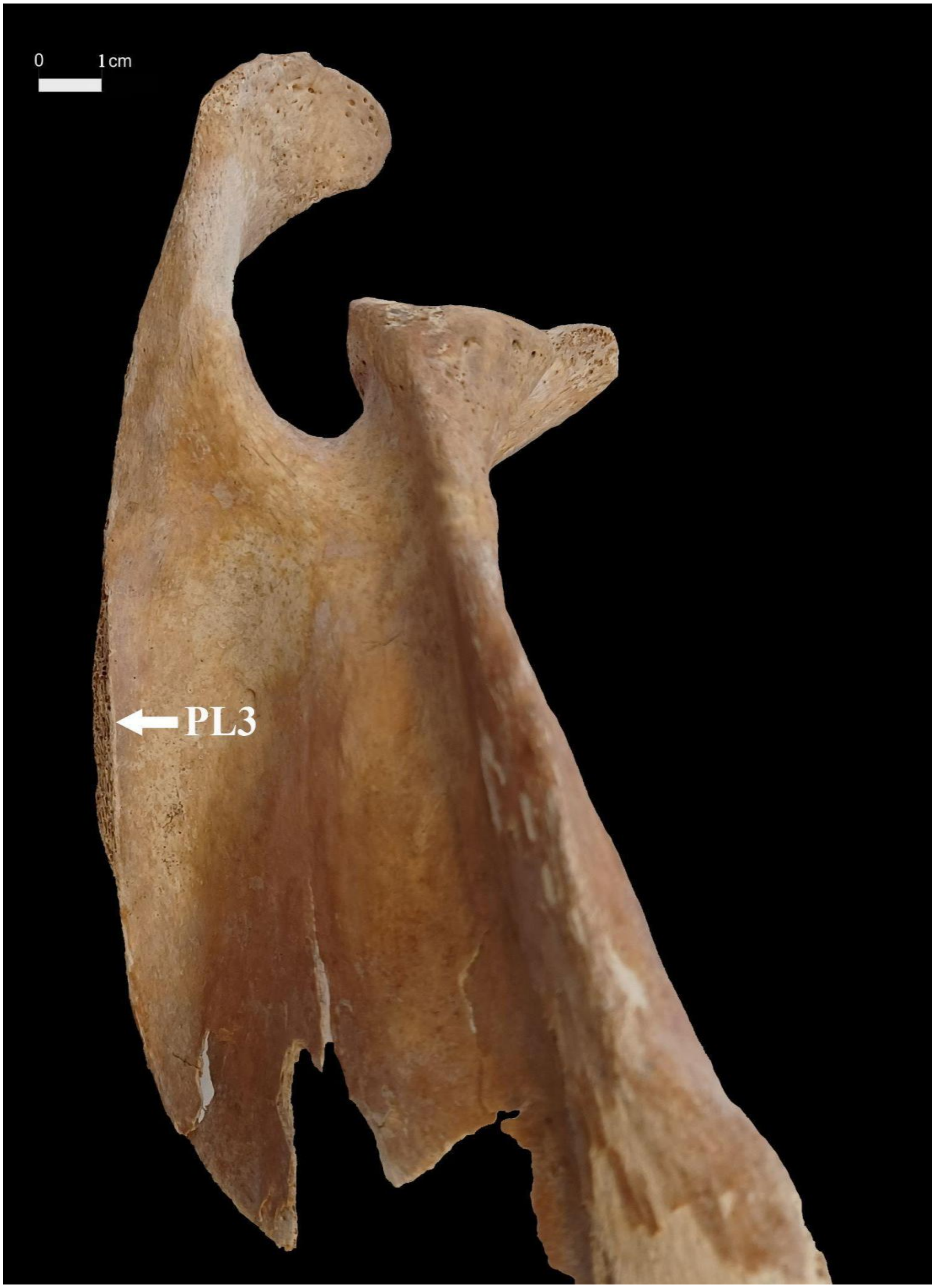


Supplementary Figure 11. A straight, tangential cut can be seen along the spine of the right scapula, with the margin cleanly clipped and a strip of bone missing. PL3= Postcranial Lesion 3. Detailed description of the perimortem lesion is available in Supplementary Information 1.2.

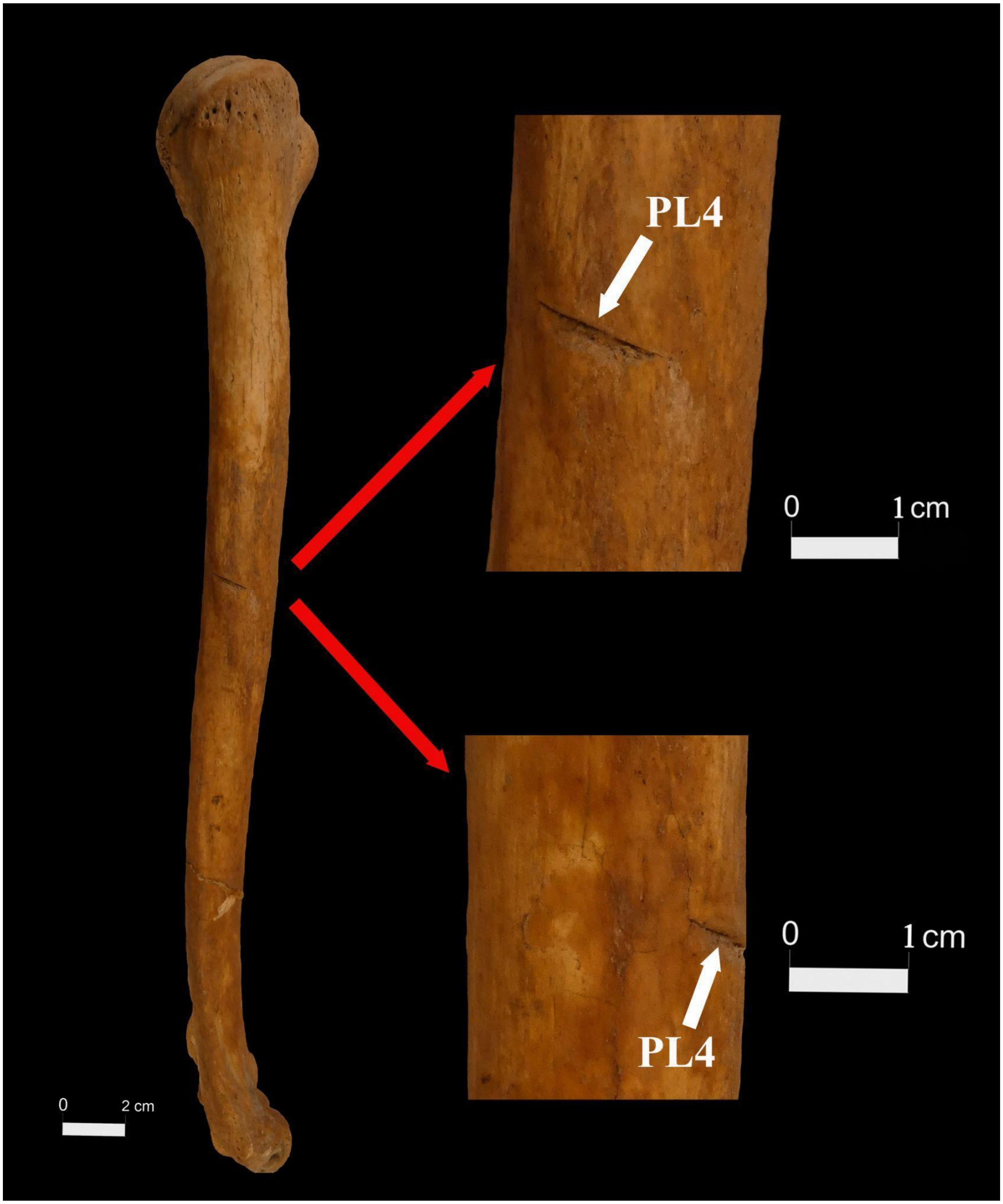


Supplementary Figure 12. A fine, superficial horizontal cut is present on the lateral-posterior surface of the tuberositas deltoidea of the right humerus. PL4= Postcranial Lesion 4. Detailed description of the perimortem lesion is available in Supplementary Information 1.2.

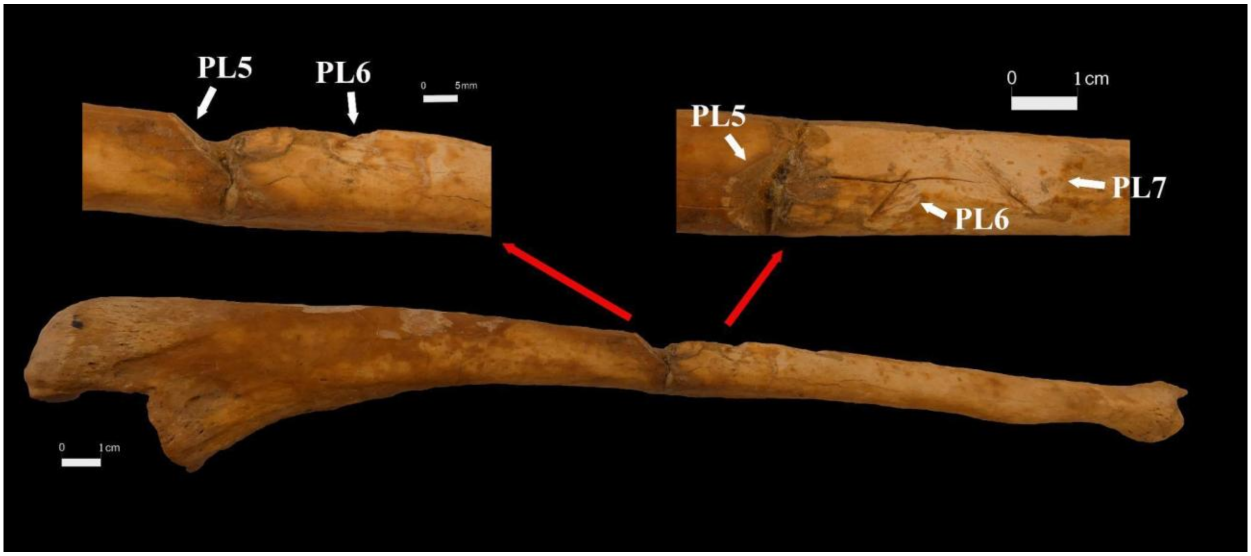


Supplementary Figure 13. Two perimortem cut marks can be observed on the right ulna. A deep, sharply angled cut is located nearly at mid-shaft on the inferior surface (PL5) and another clear cut (PL6) is present 14 mm more distal from PL5. PL5= Postcranial Lesion 5; PL6= Postcranial Lesion 6. Detailed description of the perimortem lesions is available in Supplementary Information 1.2.

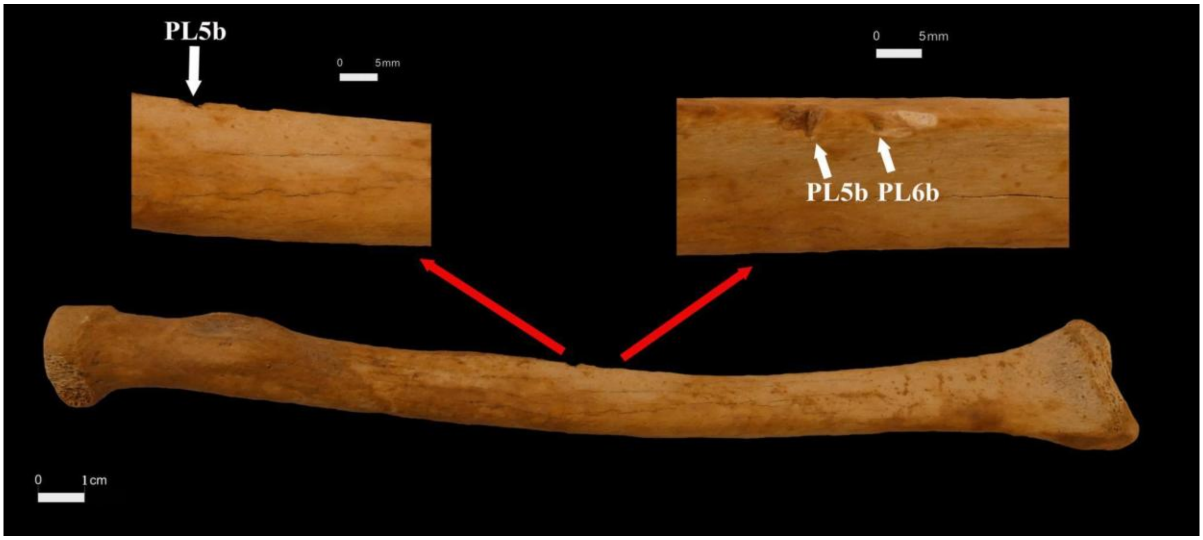


Supplementary Figure 14. Two superficial nicks on the mid-shaft medial border of the right radius (PL5b and PL6b) are likely extensions of PL5 and PL6. PL5= Postcranial Lesion 5; PL6= Postcranial Lesion 6; PL5b= Postcranial Lesion 5b;

PL6= Postcranial Lesion 6b. Detailed description of the perimortem lesions is available in Supplementary Information 1.2.

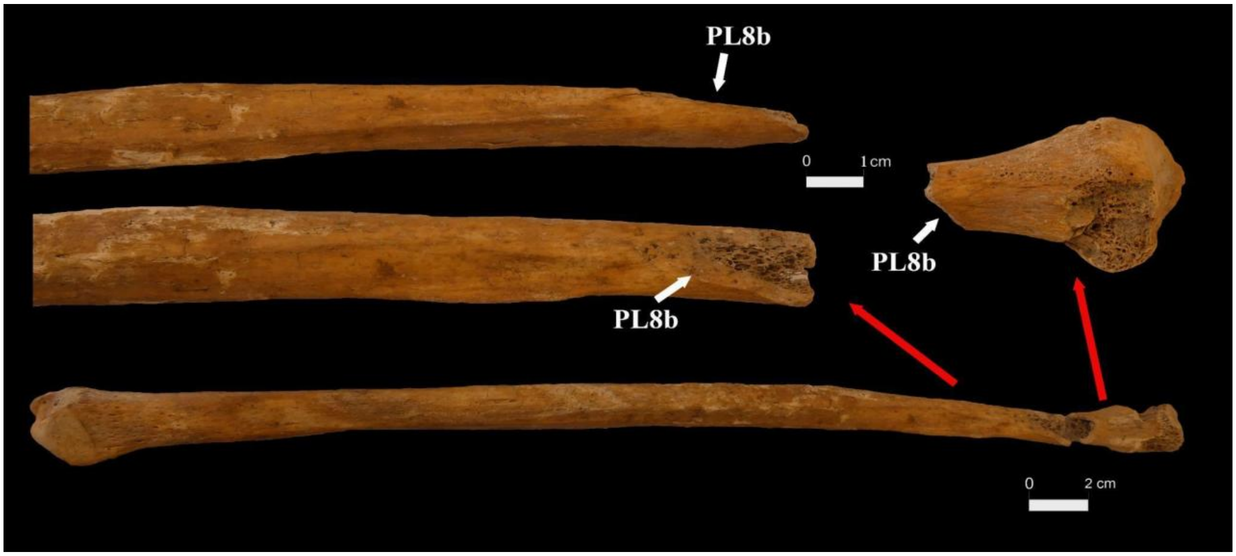


Supplementary Figure 15. A diagonal strike severing the proximal end of the left fibula. PL8b= Postcranial Lesion 8b. Detailed description of the perimortem lesion is available in Supplementary Information 1.2.

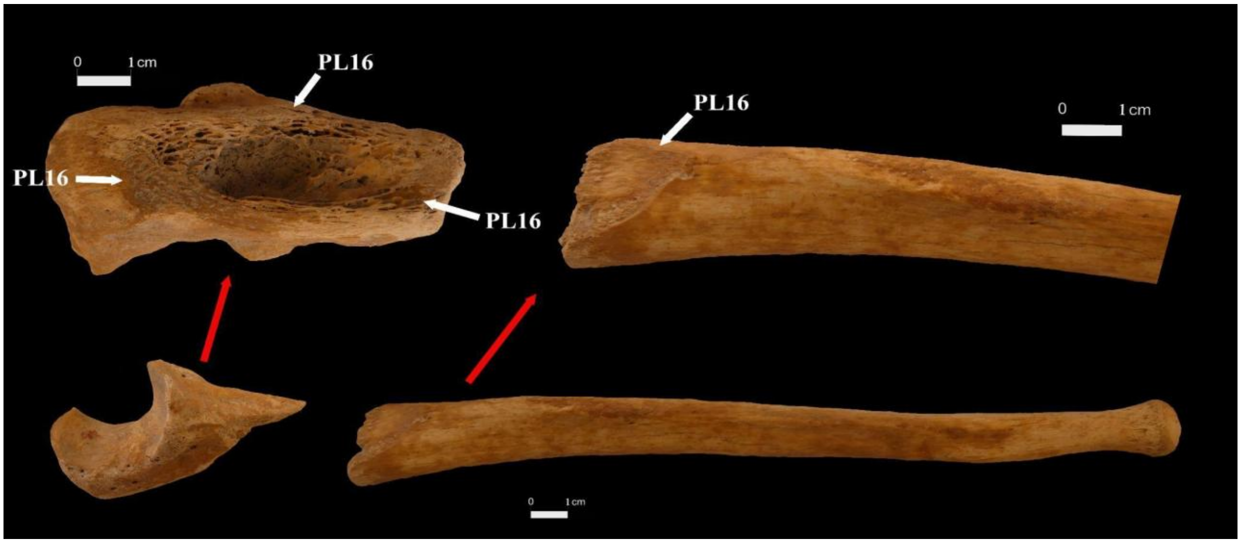


Supplementary Figure 16. A cut through the proximal end of the left ulna from the dorsal side. PL16= Postcranial Lesion 16. Detailed description of the perimortem lesion is available in Supplementary Information 1.2.

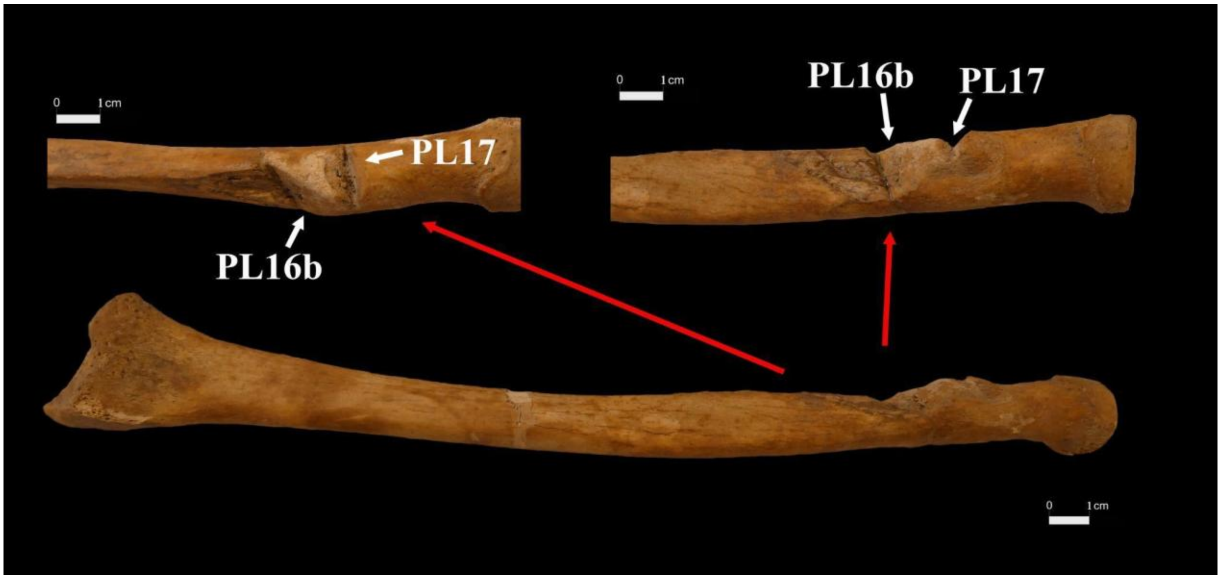


Supplementary Figure 17. Two sharp lesions cut into the proximal end of the left radius. PL16b= Postcranial Lesion 16b, PL17= Postcranial Lesion 17. Detailed description of the perimortem lesions is available in Supplementary Information 1.2.

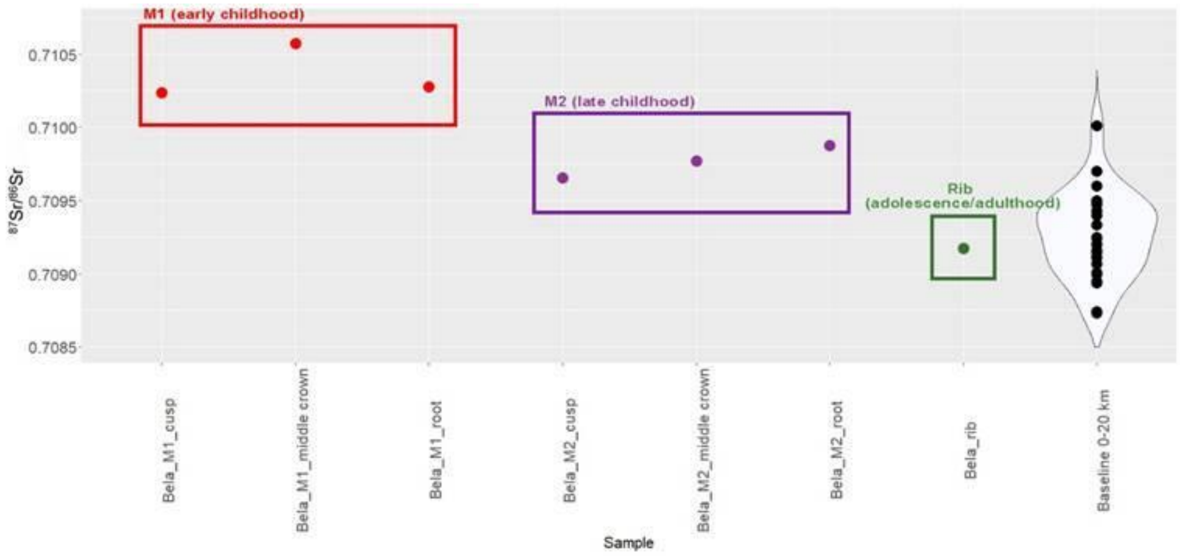


Supplementary Figure 18. $^{87}Sr/^{86}Sr$ values on different tissues of the subject, compared with local baselines, collected by Price et al. (11), Depaermentier et al. (12), Cavazzuti et al. (13)

#### Supplementary Tables (Supplementary Tables are in a separate xls file)

**Supplementary Table Legends**

Supplementary Table 1. The stable carbon and nitrogen results.

Supplementary Table 2. Y-chromosomal haplogroup determination using Yleaf

Supplementary Table 3. Sample (accession) IDs and references used to construct the mitochondrial phylogenetic tree (complete U3 dataset)

Supplementary Table 4. Details on the reference groups used in smartPCA analysis

Supplementary Table 5. Detailed results of the ancIBD analysis

Supplementary Table 6. Clinically significant nuclear variants and pigmentation-associated variants

Supplementary Table 7. Contamination predictions

#### References cited in the Supplementary Information 1-4 and in the Supplementary Tables


1 Henry AG, Hudson HF, Piperno DR. Changes in starch grain morphologies from cooking. *J. Archaeol. Sci.* **36(3)**, 915–922 (2009). https://doi.org/10.1016/j.jas.2008.11.008

2 Mickleburgh HL, Pagán-Jiménez JR. New insights into the consumption of maize and other food plants in the pre-Columbian Caribbean from starch grains trapped in human dental calculus. *J. Archaeol. Sci.* **39**, 2468–2478 (2012). https://doi.org/10.1016/j.jas.2012.02.020

3 MacKenzie L, Speller CF, Holst M, Keefe K, Radini A. Dental calculus in the industrial age: Human dental calculus in the Post-Medieval period, a case study from industrial Manchester. *Quat. Int.* **653–654**: 114–26. (2023). https://doi.org/10.1016/j.quaint.2021.09.020

4 Li Y, Zhang G, Nan P, Yang J, Cao J, Ma Z. et al. Wine or Beer? A reinvestigation of residues from bronze vessels from the Beibai'e cemetery, Shanxi China. *Heritage Science* 11: 184 (2023). https://doi.org/10.1186/s40494-023-01012-4

5 Balassa I (ed.) *Magyar Néprajz IV. Életmód.* (Akadémiai Kiadó, Budapest, 1997). http://mek.niif.hu/02100/02152/html/04/240.html

6 Radini A, Nikita E, Buckley S, Copeland L, Hardy K. Beyond food: The multiple pathways for inclusion of materials into ancient dental calculus. *Am. J. Phys. Anthropol.* **162**, 71–83 (2017). https://doi.org/10.1002/ajpa.23147

7 Power RC, Salazar-García DC, Wittig RM, Henry AG. Assessing use and suitability of scanning electron microscopy in the analysis of micro remains in dental calculus. *J. Archaeol. Sci.* **49**, 160–169 (2014). https://doi.org/10.1016/j.jas.2014.04.016

8 Horrocks, M., Nieuwoudt, M., Kinaston, R., Buckley, H., Bedford, S.Microfossil and Fourier Transform InfraRed analyses of Lapita and post-Lapita human dental calculus from Vanuatu, Southwest Pacific. *J. Roy. Soc. New Zealand* **44(1)**: 17–33 (2013). https://doi.org/10.1080/03036758.2013.842177

9 Decke, U. Mikroskopische Untersuchung an geschälten und zerkleinertenÖlsamen und Nußkernen. *Z Lebensm Unters Forch* **174**, 187–194 (1982). https://doi.org/10.1007/BF01079972

10 [Margitszigeti ásatások] [Fénykép] | Képcsarnok

11 Price TD, Knipper C, Grupe G, Smrcka V. Strontium isotopes and prehistoric human migration: the Bell Beaker period in central Europe. *Eur. J. Archaeol.* **7**: 9–40 (2024). https://doi.org/10.1177/1461957104047992

12 Depaermentier MLC, Kempf M, Bánffy E, Alt KW. Tracing Mobility Patterns through the 6th-5th Millennia BC in the Carpathian Basin with Strontium and Oxygen Stable Isotope Analyses. *PLoS One* 15(12): e0242745 (2020). https://doi.org/10.1371/journal.pone.0242745. eCollection 2020.

13 Cavazzuti C, Hajdu T, Lugli F, Sperduti A, Vicze M, Horváth A, et al. Human mobility in a Bronze Age Vatya 'urnfield' and the life history of a high-status woman. *PLoS One* **16**, e0254360 (2021). https://doi.org/10.1371/journal.pone.0254360

14 Agranat-Tamir L, Waldman S, Martin MAS, Gokhman D, Mishol N, Eshel T. et al. The Genomic History of the Bronze Age Southern Levant. *Cell* **181(5)**, 1–146-1157.e11 (2020). https://doi.org/10.1016/j.cell.2020.04.024.

15 Maár K, Varga GIB, Kovács B, Schütz O, Maróti Z, Kalmár T. et al. Maternal Lineages from 10-11th Century Commoner Cemeteries of the Carpathian Basin. *Genes* **12(3)**, 460 (2021). https://doi.org/10.3390/genes12030460

16 Saag L, Laneman M, Varul L, Malve M, Valk H, Razzak MA, et al. The arrival of Siberian ancestry connecting the Eastern Baltic to Uralic speakers further East. *Curr Biol.* **29(10)**, 1701–11.e16 (2019). https://doi.org/10.1016/j.cub.2019.04.026.

17 Damgaard PB, Marchi N, Rasmussen S, Peyrot M, Renaud G, Korneliussen T. 137 ancient human genomes from across the Eurasian steppes. *Nature 557(7705)*, 369–374 (2018). https://doi.org/10.1038/s41586-018-0094-2. Erratum in: *Nature* **563(7729)**, E16 (2018). https://doi.org/10.1038/s41586-018-0488-1.

18 Margaryan, A., Lawson, D.J., Sikora, M. et al. Population genomics of the Viking world. *Nature* **585**, 390–396 (2020). https://doi.org/10.1038/s41586-020-2688-8

19 Veeramah KR, Rott A, Groß M, van Dorp L, López S, Kirsanow K, et al. Population genomic analysis of elongated skulls reveals extensive female-biased immigration in Early Medieval Bavaria. *Proc. Natl. Acad. Sci. U. S. A.* **115(13)**, 3494–3499 (2018). https://doi.org/10.1073/pnas.1719880115.

20 Antonio ML, Gao Z, Moots HM, Lucci M, Candilio F, Sawyer S. et al. Ancient Rome: A genetic crossroads of Europe and the Mediterranean. *Science 366(6466)*, 708–714 (2019). https://doi.org/10.1126/science.aay6826.

21 Lazaridis I, Alpaslan-Roodenberg S, Acar A, Açıkkol A, Agelarakis A, Aghikyan L. et al. A genetic probe into the ancient and medieval history of Southern Europe and West Asia. *Science* **377(6609)**, 940–951 (2022). https://doi.org/10.1126/science.abq0755.

22 Maróti Z, Neparáczki E, Schütz O, Maár K, Varga GIB, Kovács B, et al. The genetic origin of Huns, Avars,

and conquering Hungarians. *Curr Biol* **32(13)**, 2858–70.e7 (2022). https://doi.org/10.1016/j.cub.2022.04.093.

23 Gerber D, Csáky V, Szeifert B, Borbély N, Jakab K, Mező Gy. et al. Ancient genomes reveal Avar-Hungarian transformations in the 9th-10th centuries CE Carpathian Basin. *Sci. Adv.* 10:eadq5864 (2024). https://doi.org/10.1126/sciadv.adq5864
